# Large-scale genetic screens identify BET-1 as a cytoskeleton regulator promoting actin health and lifespan

**DOI:** 10.1101/2022.06.22.497249

**Authors:** Gilberto Garcia, Raz Bar-Ziv, Naibedya Dutta, Darius Moaddeli, Maxim Averbukh, Toni Castro Torres, Athena Alcala, C. Kimberly Tsui, Erica A. Moehle, Ophir Shalem, Max A. Thorwald, Ryo Higuchi-Sanabria

## Abstract

The actin cytoskeleton is a three-dimensional scaffold of proteins that is a regulatory, energy-consuming network with dynamic properties to shape the structure and function of the cell. Proper actin function is required for many cellular pathways, including cell division, autophagy, chaperone function, endocytosis, and exocytosis. Deterioration of these processes manifests during aging and exposure to stress, which is in part due to the breakdown of the actin cytoskeleton. However, the regulatory mechanisms involved in preservation of cytoskeletal form and function are not well understood. Here, we performed a multi-pronged, cross-organismal screen combining a whole-genome CRISPR-Cas9 screen in human fibroblasts with *in vivo C. elegans* synthetic lethality screening. We identified the bromodomain protein, BET-1, as a key regulator of actin health and longevity. Overexpression of *bet-1* preserves actin health at late age and promotes lifespan and healthspan in *C. elegans.* These beneficial effects are mediated through actin preservation by the transcriptional regulator function of BET-1. Together, our discovery assigns a key role for BET-1 in cytoskeletal health, highlighting regulatory cellular networks promoting cytoskeletal homeostasis.

## INTRODUCTION

The actin cytoskeleton is composed of a complex network of filaments held together by actin-binding proteins. Historically, the actin cytoskeleton has been viewed as merely the structural framework of the cell, with its primary function being ascribed to cell division and the sorting and transport of cellular cargo. However, actin function is also required for many other cellular pathways, including autophagy, chaperone function, and transcriptional regulation (Balch et al., 2008; Blanpied et al., 2003; Caviston and Holzbaur, 2006; Higuchi-Sanabria et al., 2014; McCray and Taylor, 2008). The functional significance of actin in cellular health and disease is highlighted by clinical data, showing that loss of function of actin is observed in clinical manifestations, including neurodegeneration and muscle myopathies (Acsadi et al., 1991; Alim et al., 2002, 2004), and during aging (Baird et al., 2014; Higuchi-Sanabria et al., 2018; Sing et al., 2021). Specifically, actin filaments show loss of stability and marked deterioration during aging, in both single-celled yeast (Sing et al., 2021) and multiple cell-types of the multicellular nematode *C. elegans* (Higuchi-Sanabria et al., 2018), whereas changes in β-actin expression have been documented in mammals (Moshier et al., 1993). Most recently, hypotheses that unknown mechanistic pathways that can monitor and preserve the actin cytoskeleton during aging exist, have been put forward. These postulated pathway(s) would be tailored to protect the cytoskeleton, and their function might be compromised with age, contributing to decline in cellular and organismal health.

One recently identified mechanism by which the cell protects its cytoskeleton during stress is through the heat shock response (HSR), mediated by the heat shock transcription factor, HSF-1. HSF-1 is activated under thermal stress and promotes protein homeostasis through the upregulation of chaperones and other genes related to protein quality control (Morley and Morimoto, 2004). Overexpression of *hsf-1* is sufficient to confer an increase in thermal stress tolerance and lifespan in *C. elegans*, and alleviates the toxic effects associated with aging (Morley and Morimoto, 2004). Interestingly, overexpression of a truncated variant of *hsf-1* lacking the capacity to upregulate heat-shock chaperones was also sufficient to extend lifespan (Baird et al., 2014). Overexpression of this truncated form of *hsf-1* upregulated genes involved in the maintenance of the actin cytoskeleton, including the troponin C/calmodulin homolog, *pat-10*, which is both sufficient and necessary for HSF-1-mediated thermotolerance and longevity (Baird et al., 2014). However, many actin-regulating genes are not bona fide targets of HSF-1, suggesting the existence of other master regulators, which function independently or in parallel with HSF-1 to modulate actin and protect the cytoskeleton. For example, the general regulation of actin is governed by key actin nucleation factors that fall into three major classes 1) the Arp2/3 (actin-related protein 2/3) complex that builds branched actin networks (Machesky et al., 1994, 1999); 2) formins that build unbranched actin filaments (Pruyne et al., 2002); and 3) tandem-monomer binding nucleators, which bring monomers together to form actin nucleation seeds (Quinlan et al., 2005; Sagot et al., 2002). All three types of actin nucleation factors were not found to be regulated by HSF-1, and the beneficial effect of *hsf-1* overexpression was actually independent of the tropomyosin, *lev-11* (LEVamisole resistant) (Baird et al., 2014).

To identify previously unidentified regulators of actin health, we adopted a multi-pronged, cross-organismal screening approach. Since actin is one of the most highly conserved proteins, both in terms of sequence and functional homology, we held the rationale that an evolutionary conserved perspective could provide a powerful method to identify key regulators of actin function. We first utilized a CRISPR/Cas9-driven growth-based genetic screen in human cells to identify genes required for survival under actin stress. Actin stress was applied by exposure to cytochalasin D, a drug that inhibits actin polymerization by binding to F-actin filaments and preventing the addition of actin monomers (May et al., 1998). We then performed a secondary screen of the top ∼500 candidate genes in the multicellular nematode, *C. elegans*, allowing us not only to explore its cellular contributions, but also to associate physiological manifestations in a multicellular organism. Our secondary screen consisted of a synthetic lethality screen to identify genes that when knocked down cause lethality in animals exposed to a sublethal knockdown of actin. These screens identified *bet-1* (two bromodomain family protein), which promotes actin health, longevity, and healthspan in a multicellular eukaryote.

## RESULTS

In our work to identify genes critical for actin health, we chose to inhibit actin polymerization using the chemical drug cytochalasin D. Cytochalasin D prevents actin polymerization by binding to F-actin filaments and blocking the addition of actin monomers (May et al., 1998). We screened for a concentration of cytochalasin D that only mildly affected growth rate (Fig. S1A) and performed a CRISPR/Cas9-driven reverse genetic screen in BJ fibroblasts, a karyotypically normal human fibroblast line. Next, we performed a whole-genome CRISPR knockout screen using the AVANA pooled sgRNA library (Schinzel et al., 2019; Shalem et al., 2014) in cells treated with 0.1 µM cytochalasin D (Fig. 1A). We compared gene-based depletion p-values between control and treatment arms to curate a list of genes that were enriched or depleted in our cytochalasin-treated cells (**Table S1-2**, Fig. S1B). Importantly, one of the most significantly depleted genes was the actin-encoding gene, ACTB. Moreover, gene ontology (GO) analysis (Chen et al., 2013; Kuleshov et al., 2016; Xie et al., 2021) revealed actin cytoskeleton, the general cytoskeleton, and actin filaments as significantly enriched GO terms for the top depleted genes (Fig. S1C). These data provide confidence that our screen is revealing genes important for actin function. Interestingly, many mitochondrial genes were found amongst significantly enriched genes (Fig. S1D), though this is consistent with many findings that have previously revealed actin-mitochondrial interactions (Fehrenbacher et al., 2004; Higuchi et al., 2013; Moehle et al., 2021; Tharp et al., 2021).

**Fig. 1.**
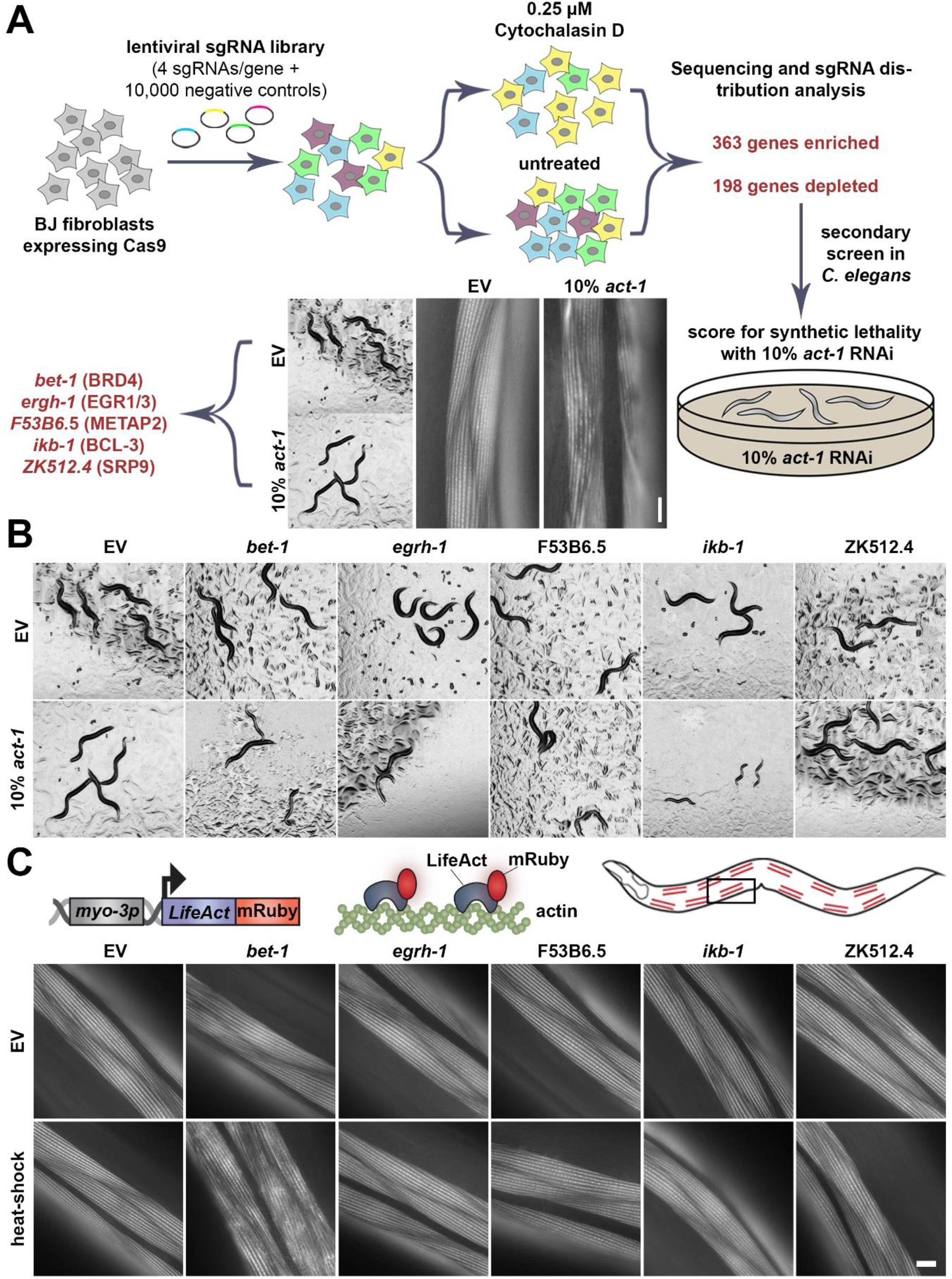
Multiplex screening reveals *bet-1* is involved in regulation of actin. **(A)** Schematic of cross-species screening. CRISPR-Cas9 screening was performed in BJ fibroblasts with 0.1 µM cytochalasin D and top hits were screened in *C. elegans* with 90% candidate RNAi and 10% *act-1* RNAi. **(B)** Images of N2 animals grown on 90% candidate RNAi mixed with either 10% empty vector (EV) or 10% *act-1*. Hits were defined as those that exhibit an observable phenotype when combined with *act-1* RNAi, but not with EV. **(C)** LifeAct::mRuby is expressed specifically in the body wall muscle cells with the *myo-3* promoter. LifeAct::mRuby binds to F-actin filaments to allow visualization of actin. Representative fluorescent images of body wall muscle actin are shown. Animals were grown on RNAi from hatch. Animals were heat-shocked for 1 hr at 37 °C and immediately imaged. Images were captured on a Zeiss AxioObserver.Z1. Scale bar is 10 µm.

To follow up on the identified hits and link their contribution to cellular and organismal physiology, we opted for performing a cross-species analysis in the nematode *C. elegans* with the rationale that genes important for surviving actin stress in two species must be critical regulators of actin health and function. Indeed, we found in a previous study that similar cross-species screening approaches provided more biologically meaningful data than computational methods alone to identify candidate genes that impact lifespan (Moehle et al., 2021). Therefore, we searched the top ∼500 enriched and depleted genes from our screen for evolutionary-conserved orthologs in *C. elegans* (Kim et al., 2018) and identified a list of ∼400 genes. Next, we performed a synthetic lethality screen for these candidate genes, searching for genes that when knocked down, caused lethality when combined with a sub-lethal knockdown of actin (*act-1*) (Fig. 1A). Importantly, the selected RNAi sequence of *act-1* shows overlap with all 5 isoforms of actin in *C. elegans* (*act-1* through *act-5*) and exhibits visible perturbations of actin health without affecting whole organismal physiology. Animals treated with 10% *act-1* RNAi develop normally to adulthood but display notable perturbations of actin quality in muscle cells (Fig. 1A). Therefore, we proceeded with our synthetic lethality screen by performing RNAi knockdown of candidate genes mixed with *act-1* RNAi in a 9:1 ratio, respectively.

Our secondary screen revealed 5 potential candidate genes with varying phenotypes (Fig. 1B). RNAi knockdown of *bet-1*, *egrh-1* (Early Growth factor Response factor Homolog), *F53B6.5*, and *ikb-1* (I Kappa B homolog) resulted in delayed development when combined with 10% *act-1* RNAi but exhibit no physiological phenotypes when knocked down alone. RNAi knockdown of *ZK512.4* did not exhibit a developmental defect but showed sterility (no eggs were visible on the plate at day 1 of adulthood) when combined with 10% *act-1* RNAi, despite showing visible progeny formation when knocked down alone. *ZK512.4* is a predicted ortholog of human SRP9 (signal recognition particle), a critical part of protein targeting to the endoplasmic reticulum (ER) (Mary et al., 2010), with no known association with the cytoskeleton, though studies have shown that actin can impact ER dynamics (Korobova et al., 2013; Poteryaev et al., 2005). *egrh-1* is the ortholog of human EGR1 (early growth response 1), a major transcription factor that has been implicated in multiple diseases including cancer (Wang et al., 2021), neuropsychiatric disorders (Duclot and Kabbaj, 2017), and Alzheimer’s disease (Qin et al., 2016). *F53B6.5* is a predicted ortholog of human methionyl aminopeptidase 2 (METAP2), which removes methionine residues from nascent polypeptide chains and are also implicated in cancer (Yin et al., 2012). *bet-1* is the ortholog of human BRD4, a bromodomain protein involved in cell fate decisions in both *C. elegans* (Shibata et al., 2010) and mammals (Lee et al., 2017; Linares-Saldana et al., 2021), and also implicated in cancer progression (Huang et al., 2016).

Next, we sought to determine which of our candidate genes directly impacted actin integrity and lifespan. To directly measure actin organization, we used animals expressing LifeAct::mRuby in the muscle. Here, the F-actin binding protein, LifeAct is fused to a fluorescent molecule to allow robust visualization of actin integrity in muscle cells (Higuchi-Sanabria et al., 2018). RNAi knockdown of *bet-1* consistently resulted in decreased lifespan, whereas RNAi knockdown of *egrh-1, F53B6.5,* and *ikb-1* resulted in inconsistent changes to lifespan (Fig. S2A, **Table S3-S4**). Surprisingly, RNAi knockdown of any of the five candidate genes had no impact on actin integrity (Fig. 1C**, top**). However, our previous work showed that RNAi knockdown of *hsf-1*, which has dramatic effects on organismal health and lifespan (Hsu et al., 2003; Morley and Morimoto, 2004), also did not impact actin organization in the muscle in young, unstressed adults, and instead only exhibited measurable phenotypes during stress or aging (Higuchi-Sanabria et al., 2018). Therefore, we applied an acute exposure to heat stress (1 hr at 37 °C), which has no measurable effect on wild-type animals. However, RNAi knockdown of *bet-1* – and no other candidate genes – resulted in a dramatic loss of actin integrity and organization under acute heat stress (Fig. 1C**, bottom**). Therefore, as a gene that directly impacts actin integrity and aging, we pursued *bet-1* for follow-up analysis.

Our data suggest that loss of *bet-1* results in decreased stability of actin, such that acute exposure to stress is sufficient to perturb actin function. In addition, we find that knockdown of *bet-1* resulted in the premature breakdown of actin during aging, such that muscle actin showed marked deterioration as early as day 4, compared to wild-type animals that only start to show mild signs of actin disorganization between day 7-10 (Fig. 2A). The RNAi knockdown of *bet-1*, as validated by reduced transcript levels using RT-PCR (Fig. S2B), also caused a decrease in lifespan (Fig. 2B). Moreover, a *bet-1* mutant generated by introducing a premature stop codon at amino acid 17, consistently showed a significant decrease in lifespan (Fig. S2C). Surprisingly, overexpression of the primary isoform of *bet-1*, *bet-1A,* previously shown to be required for its canonical function in maintenance of cell fate (Shibata et al., 2010) had no impact on lifespan. Similarly, overexpression of the alternative isoform *bet-1C* did not impact lifespan. Instead, overexpression of the *bet-1* isoform *bet-1B* significantly increased lifespan (Fig. 2C), which was reversed by *bet-1* RNAi, suggesting that this is not due to an off-target effect (Fig. 2B). Importantly, overexpression of *bet-1B* also protected actin filaments at advanced age, as animals with *bet-1B* overexpression display no measurable changes to actin filaments as late as day 13 when wild-type animals show obvious collapse (Fig. 2A). This increase in actin filament stability also results in increased muscle function, which can be measured by increased motility both at young and old age (Fig. 4B, see blue and green), in agreement with a previous study, which described a premature loss of motility in *bet-1* loss of function animals (Fisher et al., 2013). These data provide direct evidence that *bet-1* is a bona fide regulator of actin function, which has direct implications in organismal health and longevity.

**Fig. 2.**
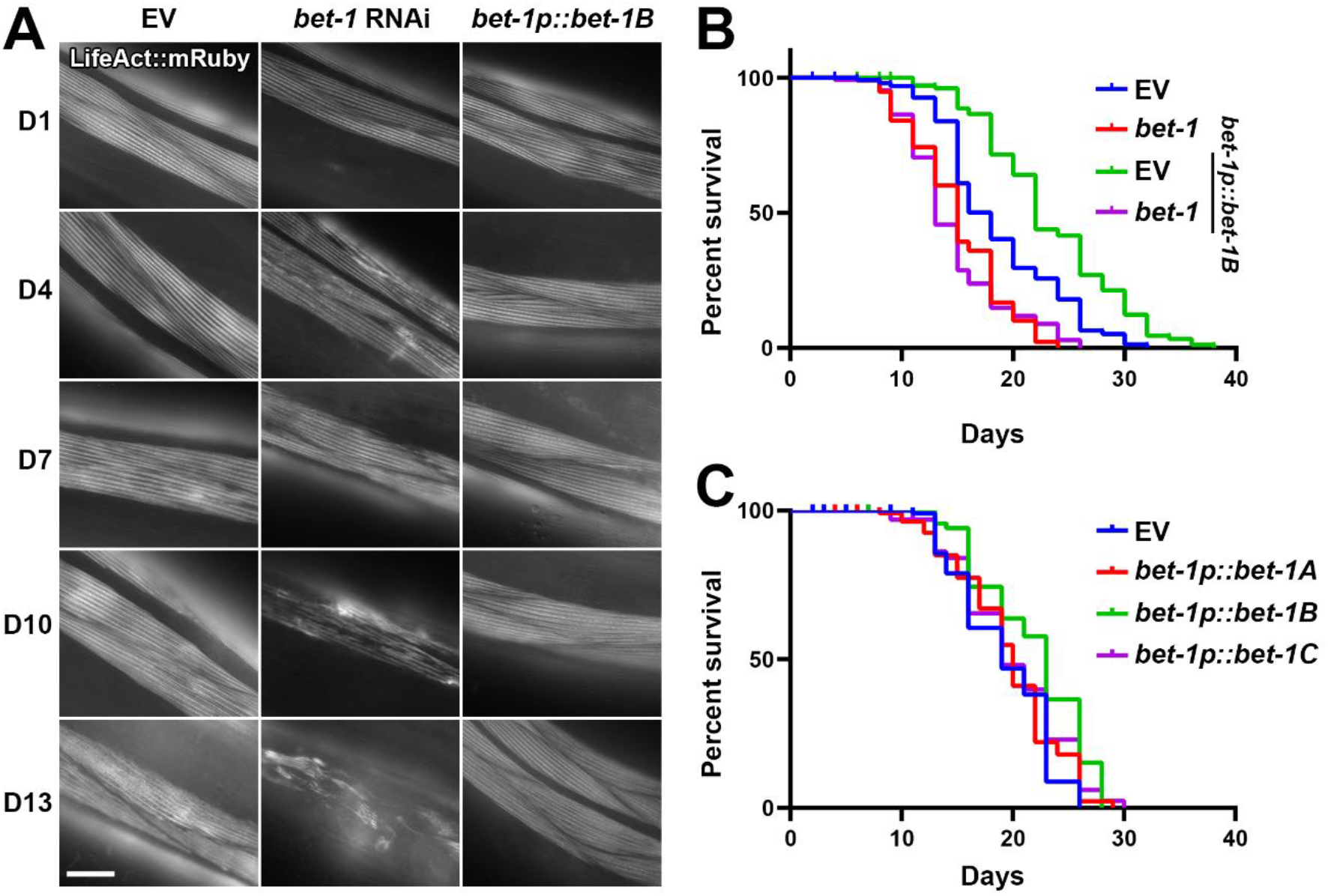
*bet-1* expression directly impacts actin health and lifespan. **(A)** Representative fluorescent images of adult animals expressing Lifeact::mRuby from a muscle-specific promoter *myo-3p*. N2, wild-type animals were grown on empty vector (EV) or *bet-1* RNAi from hatch, and *bet-1B* overexpression animals (*bet-1p::bet-1B*) were grown on EV from hatch. All animals were imaged at day 1, 4, 7, 10, and 13 of adulthood. Images were captured on a Zeiss AxioObserver.Z1. Scale bar is 10 µm. **(B)** Lifespans of N2 (blue) and *bet-1B* overexpression (green) animals grown on EV or *bet-1* RNAi (N2, red; *bet-1B* overexpression, purple) from hatch. See **Table S3-4** for lifespan statistics. **(C)** Lifespans of N2 (EV, blue) and overexpression of *bet-1A* (*bet-1p::bet-1A*, red), *bet-1B* (*bet-1p::bet-1B*, green), and *bet-1C* (*bet-1p::bet-1C*, purple) grown on EV from hatch. See **Table S3-4** for lifespan statistics.

BET-1 is a conserved double bromodomain protein that functions as a transcriptional regulator to maintain stable cell fates (Shibata et al., 2010). Specifically, it recognizes lysines of histone tails acetylated by the MYST family of acetyltransferases (MYST HATs), and loss of *mys-1* and *mys-2* results in mislocalization of BET-1 and loss of BET-1 function (Shibata et al., 2010). To determine whether the functional role of BET-1 in aging and cytoskeletal maintenance were similar to those of cell fate decisions, we first synthesized a GFP-tagged variant of BET-1B. Indeed, GFP::BET-1B also localize primarily to the nucleus and creates distinct punctae, consistent with previous studies (Fig. S3A) (Shibata et al., 2010). In contrast to cell fate decisions, *mys-1* RNAi was not sufficient to fully phenocopy loss of *bet-1*, as animals did not exhibit premature deterioration of actin organization despite having a comparable decrease in lifespan, which were not additive with *bet-1* knockdown (Fig. S3B, S4A). This may be due to an incomplete knockdown of *mys-1* via RNAi (Fig. S2B). However, RNAi knockdown of *mys-1* fully suppressed the lifespan extension (Fig. S4A) and protection of the cytoskeleton at late age found in *bet-1B* overexpression animals (Fig. S4B). Taken together, our data suggest that although some differences exist, similar to cell fate decisions, BET-1 promotes actin integrity and lifespan downstream of MYST HATs, likely as a transcriptional regulator.

Thus, to directly test a functional role for BET-1 in transcriptional regulation of cytoskeletal gene programs, we performed RNA-seq in *bet-1* loss of function and *bet-1B* overexpression animals. We used both RNAi knockdown and the newly generated *bet-1* mutant. As expected, we observed a high correlation between the expression profiles of the knockdown and knockout of *bet-1* (Fig. S5A-C). We focused our analysis on the *bet-1B* overexpression (Fig. 3A) and found that genes associated with the general cytoskeleton and the actin cytoskeleton were among the most enriched gene ontology terms (Fig. 3B), similar to enrichments observed in the initial human fibroblast screen (Fig. S1). Other enriched terms included microtubules, chromatin, and vesicle-associated terms. Congruent with our observations from our cytoskeletal imaging data, these data suggest that BET-1 mediated longevity was through its effects on the cytoskeleton. Therefore, we further focused on all genes related to actin function as previously annotated (Holdorf et al., 2020). We found that out of the five actin genes in *C. elegans*, the expression of *act-3* was significantly induced. Furthermore, we observed additional changes, including induction and repression, of genes involved in the actin cytoskeleton (Fig. 3C), with many opposing effects between the loss of function and *bet-1B* overexpression datasets across different cytoskeleton-related gene groups (Fig. S5D). Strikingly, these transcriptional changes were dependent on *mys-1,* as *mys-1* knockdown reversed most of the induction and repression of differentially expressed genes, including the induction in *act-3* (Fig. S5E-F). These data are consistent with a model whereby BET-1 impacts lifespan and actin health through its role as a transcriptional regulator downstream of MYS-1.

**Fig. 3.**
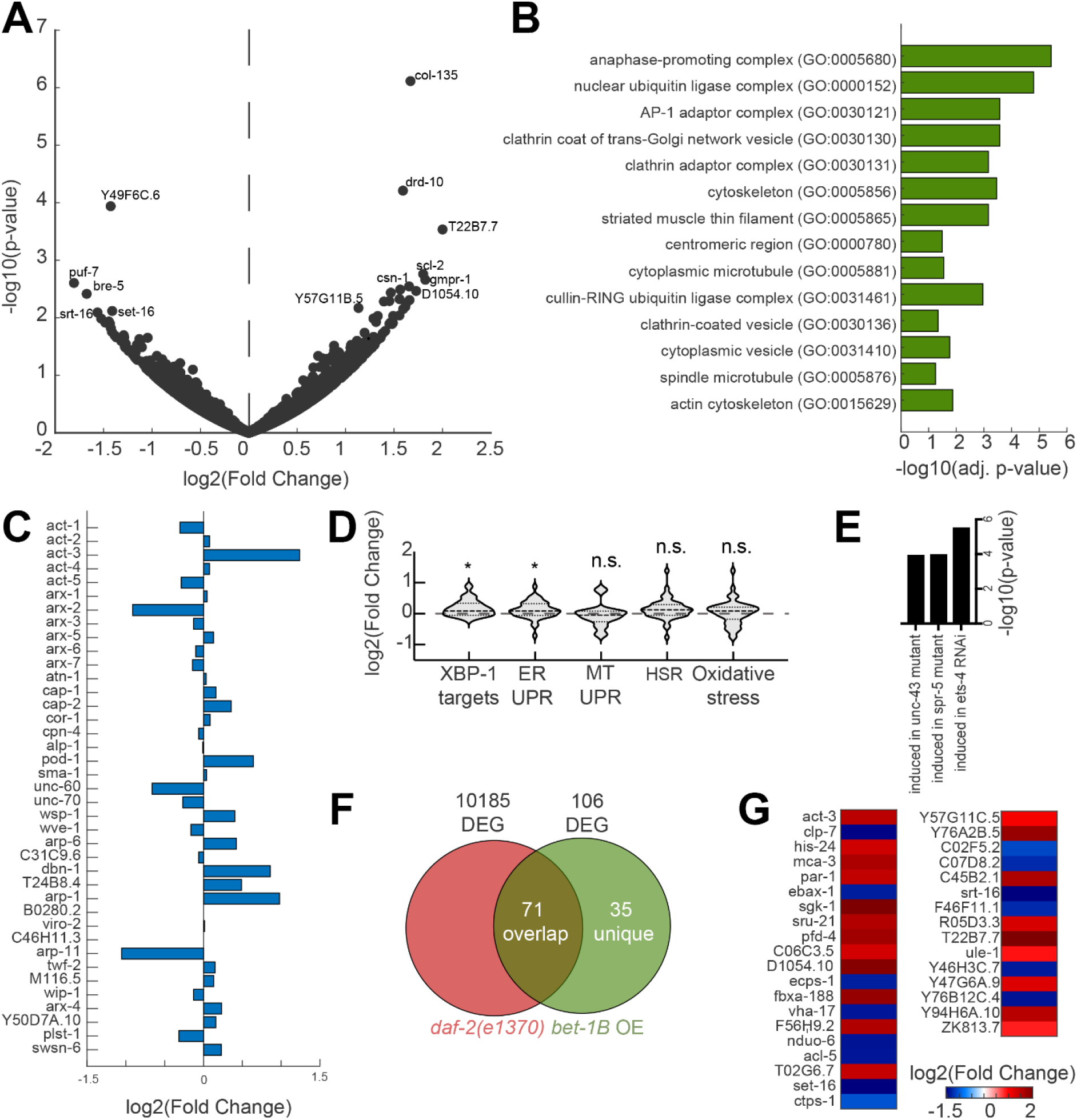
Overexpression of *bet-1* overexpression drives changes in cytoskeletal regulatory genes. **(A)** Differentially expressed genes upon overexpression of *bet-1B*: Volcano plot of the genome-wide changes in gene expression upon over-expressing bet-1B, as compared to an N2 wildtype control. **(B)** Gene ontology enrichments (Chen et al., 2013) for differentially expressed genes (*p-value* < 0.05) in worms over-expressing *bet-1B*. **(C)** log2(Fold changes) for all genes annotated as cytoskeleton: actin function in WormCat (Holdorf et al., 2020). **(D)** Gene expression changes in groups of genes linked to longevity: UPR^ER^ GO:0030968; Mitochondrial UPR (UPR^MT^) GO:0034514; Heat shock response (HSR) GO:0009408; Oxidative stress response GO: 0006979. XBP-1 targets as previously defined (Urano et al., 2002). **(E)** Comparison of differentially expressed genes (*p-value* < 0.05) in *bet-1B* over-expressing worms as compared to a long-lived *daf-2(e1370)* mutant (Zarse et al., 2012) reveals a group of 35 unique genes, plotted in **(F)**. **(G)** WormExp (Yang et al., 2016) analysis integrating previously published datasets reveals significant enrichment for the annotated perturbations. See **Table S5** for all differentially expressed genes.

We then tested whether other protective pathways that have been linked to longevity were altered upon *bet-1B* activation. We could not find activation of the mitochondrial unfolded protein response (UPR^MT^), the heat shock response (HSR), or oxidative stress response. We did observe a mild activation of genes involved in the endoplasmic reticulum UPR (UPR^ER^). These data were further verified by direct comparison to a dataset identifying targets of the UPR^ER^ transcription factor, XBP-1 (Fig. 3D). We also compared our dataset to a previously published gene expression dataset of *daf-2(e1370)* (Zarse et al., 2012), a mutant with a reduced function of the insulin/IGF-1 receptor signaling, which has been implicated in aging. We found that 71 out of 106 differentially expressed genes overlapped between worms overexpressing *bet-1B* and the long-lived *daf-2(e1370)* (Fig. 3E). Among the genes unique to the *bet-1B* overexpressing worms was *act-3* (Fig. 3F). Interestingly, our investigation of genes that were significantly induced in the *bet-1B* overexpressing worms showed a similar transcriptional signature to that induced by knockdown of the transcription factor *ets-4/SPDEF* (erythroblast transformation specific/SAM pointed domain containing ETS transcription factor), which has been previously linked to aging (Thyagarajan et al., 2010), potentially through regulation of the actin cytoskeleton via VASP (vasodilator-stimulated phosphoprotein) (Bear et al., 2002; Ye et al., 2020) (Fig. 3G). In addition, we also observed a similarity with genes affected by knockdown of the methyl demethylase *spr-5* (suppressor of presenilin), another gene whose reduction has been linked to longevity (Greer et al., 2016). Together, these data provide further evidence that the pro-longevity transcriptional program induced by *bet-1B* is at least in part due to a dedicated program to promote cytoskeletal health.

To experimentally validate our findings from transcriptome analysis, we tested the impact of destabilizing the actin cytoskeleton on the beneficial effects of *bet-1B* overexpression. Importantly, perturbations of actin function completely suppressed the lifespan extension found in *bet-1B* overexpression animals (Fig. 4A). Specifically, actin function was perturbed by 10% *act-1* RNAi, similar to conditions used for our synthetic lethality screens. 10% *act-1* RNAi results in a significant decrease in lifespan extension, and overexpression of *bet-1B* has no impact on lifespan in these animals. Thus, the lifespan extension of *bet-1B* overexpression is indeed dependent on its alterations of the actin cytoskeleton, as knockdown of actin was sufficient to reverse the longevity phenotype. Similarly, perturbation of actin function suppressed the beneficial effects of *bet-1B* overexpression on motility (Fig. 4B). These data suggest that the beneficial effects of *bet-1B* are primarily due to its role in promoting cytoskeletal function, likely as a transcriptional regulator.

**Fig. 4.**
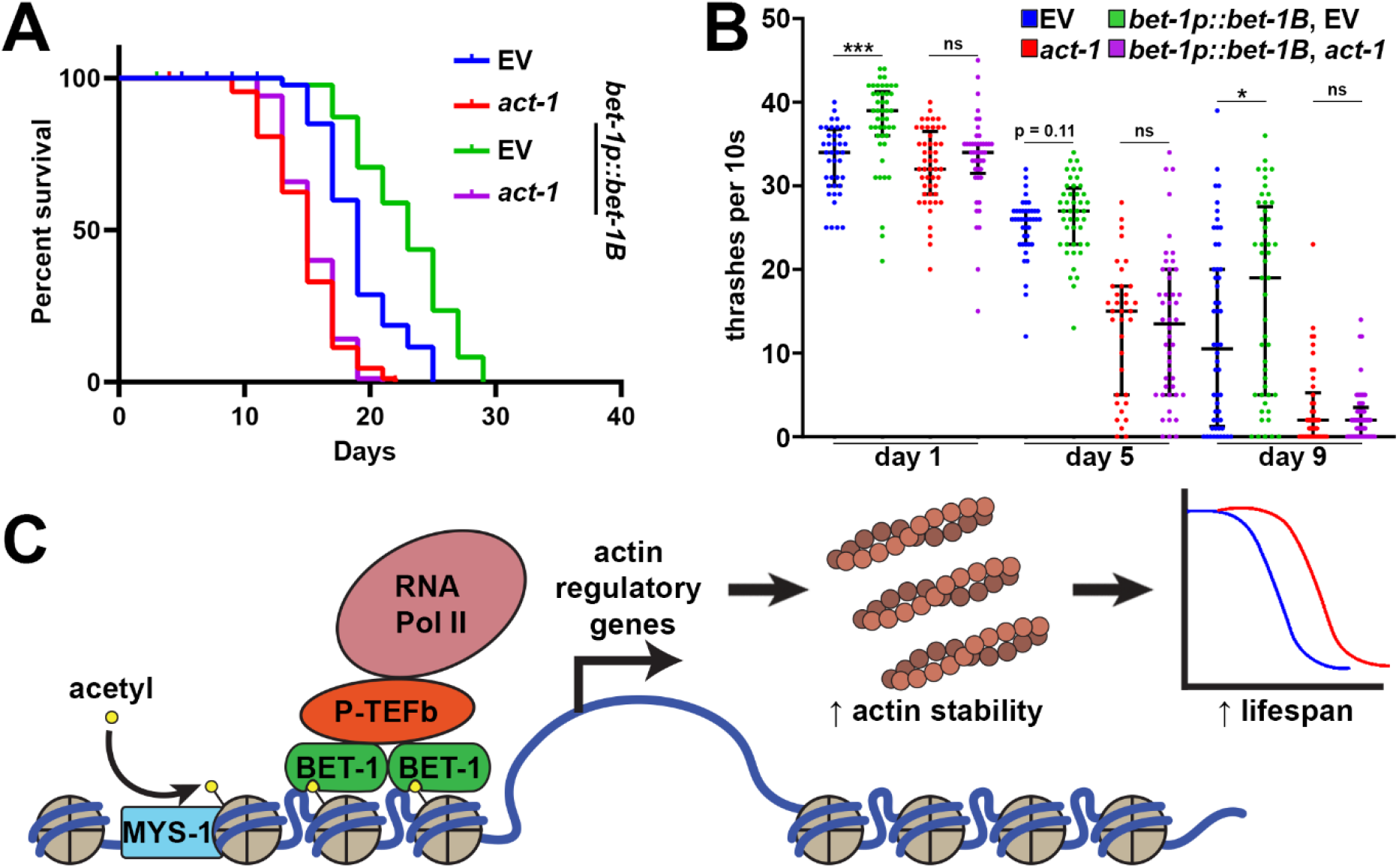
*bet-1* overexpression promotes lifespan and healthspan through effects on actin. **(A)** Lifespans of wild-type, N2 (EV, blue) and *bet-1B* overexpression (*bet-1p::bet-1B*, green) animals grown on EV or *10% act-1* RNAi (N2, red; *bet-1B* overexpression, purple) from hatch. See **Table S3-4** for lifespan statistics. **(B)** Thrashing assays were performed on N2 (EV, blue) and *bet-1B* overexpression (*bet-1p::bet-1B*, green) animals grown on EV or *10% act-1* RNAi (N2, red; *bet-1B* overexpression, purple) from hatch. Animals were grown on FUDR to prevent progeny development and assayed on day 1, 5, and 9 of adulthood. Recordings were performed in M9 solution on a Leica M205FCA stereomicroscope with a Leica K5 camera and thrashing was scored manually over a 10 second recording. Data is representative of three independent trials. n = 36-54 worms per condition. *** = p < 0.001; * = p < 0.05; ns = p > 0.05 calculated using non-parametric Mann-Whitney testing. Each dot represents a single animal and lines represent median and interquartile range. **(C)** Model for BET-1 regulation of actin. MYS-1 acetylates histones (Ceol and Horvitz, 2004), allowing recruitment of BET-1 to chromatin (Shibata et al., 2010). BET-1 recruitment to chromatin allows for recruitment of transcription initiation factors like P-TFEb, which allows recruitment of RNA Pol II (Jang et al., 2005) to promote transcription of actin regulatory genes, which promotes actin function and lifespan.

## DISCUSSION

The actin cytoskeleton is a complex and dynamic cellular organelle, which requires a tight regulation of the biosynthesis of its building blocks, their polymerization, disassembly and breakdown. The dynamic rearrangements of actin underlie many cellular functions, and have critical implications on organismal phenomena, such as disease states and overall health. Many previous works focused on understanding the interactions between the players which directly take part in the cytoskeleton. For example, a competition model has been proposed to exist between actin assembly factors and monomeric actin (Davidson and Wood, 2016). Inspired by the protein homeostasis pathways of other organelles (Dutta et al., 2022), we were intrigued to explore whether a master transcriptional regulator may exist, which drives and promotes cytoskeletal health. Interestingly, our cross-species screen revealed potentially evolutionarily conserved regulators of actin homeostasis between mammals and *C. elegans*. Through CRISPR-Cas9 screening in karyotypically normal human fibroblasts, we identified genes that when knocked out impact survival under actin destabilization caused by exposure to cytochalasin. Cytochalasin D is a mycotoxin that inhibits actin polymerization by binding to F-actin and preventing polymerization of actin monomers (May et al., 1998). Gene ontology analysis of our significantly depleted genes identified actin cytoskeletal organization as the highest enriched gene set, as well as genes involved in cell-matrix adhesion, cell motility, wound healing, intracellular transport, and autophagy. These pathways all mechanistically require a properly functional actin cytoskeleton, which gave us high confidence in our initial screen. However, a few limitations of our screen are that we used the Avana sgRNA KO library, which since its production has been outperformed by some other libraries, especially CRISPR interference (CRISPRi) or CRISPR activation (CRISPRa) platforms, which can enable more flexible gene modulation (Sanson et al., 2018). In addition, our utilization of cytochalasin D can be limiting due to the wide-range effects of the drug (Foissner and Wasteneys, 2007).

Rather than optimize our primary screen with alternative libraries or other actin-destabilizing drugs (e.g., latrunculin), we opted for a secondary screening platform in *C. elegans*. We reasoned that as one of the most highly conserved gene/protein across eukaryotes in sequence and function, a cross-species approach would be optimal to identify critical regulators of actin. In addition, *C. elegans* are an *in vivo* whole animal model, free from some of the restrictions or caveats of an *in vitro* cell culture system, such as major changes to actin cytoskeletal integrity when grown in a plastic dish (Tharp et al., 2021). Finally, phenotypic analysis in *C. elegans* is not limited solely to growth rate, and has multiple levels of phenotyping, including development, motility, fecundity, size, and overall physiological health, all of which can be rapidly screened simultaneously. For our secondary screen, we used a genetic screening method whereby we performed partial knockdown of all actin genes using RNAi. We opted for this method as the thick cuticle of *C. elegans* lead to low permeability of many drugs (Xiong et al., 2017), making the usage of cytochalasin D not only cost-prohibitive for this study, but low confidence in maintaining a homogenous effect. Moreover, the actin cytoskeleton is important for cuticle development in worms (Costa et al., 1997) further compounding on this problem, and thus screening for regulators of actin using a drug may introduce several caveats that can bias hits for those involved in cuticle development. Overall, our two-species screening platform attempted to make use of two robust platforms equipped with a unique set of benefits that allowed for identification of evolutionary conserved regulators of actin.

Through our cross-species approach, we identified *bet-1*, the *C. elegans* ortholog of human BRD2 and BRD4. *bet-1* encodes a double bromodomain protein that has been originally characterized for its role in cell fate decisions (Shibata et al., 2010). In addition, BET-1 has been found to directly impact muscle myosin levels during aging, likely as a transcriptional regulator (Fisher et al., 2013). Actin filaments are usually in direct association with myosin (Cooper, 2000) and actin-myosin interactions are critical for *C. elegans* body wall muscle structure and function (Gieseler et al., 2017). Moreover, BRD4 has been implicated in cancer aggression as an angiogenesis promoting factor, a process that directly involves actin polymerization (Huang et al., 2016). Interestingly, BRD2 (an alternative homolog of *bet-1*) dosage has been previously linked to longevity in C57B6/J mice, although the underlying molecular basis was not identified (Pathak et al., 2020). While these previous studies showed potential correlation of BET-1/BRD2/BRD4 with actin function and/or longevity, our study has now provided evidence that expression of *bet-1* directly impacts actin organization and function, which has direct significance in longevity. Specifically, loss of function of *bet-1* results in premature breakdown of actin during aging, while its overexpression protects actin filaments at late age and promotes both healthspan and lifespan.

Our fluorescent studies suggest that BET-1 does not impact actin through direct protein interactions, as all visible BET-1 protein is found within the nucleus and localizes to puncta which in no way resembles actin filaments. Moreover, the beneficial effects of BET-1 on actin and lifespan are dependent on MYS HATs, which acetylate histones at specific lysine residues, allowing BET-1 binding to alter gene expression (Shibata et al., 2010). Thus, BET-1 likely impacts actin function as a transcriptional regulator that promotes expression of actin regulatory genes (Fig. 4C). Indeed, our transcriptome analysis revealed that cytoskeletal regulators make up a large majority of upregulated genes in *bet-1* overexpressing animals, with *act-3* specifically being highly induced. We also observed changes in other cytoskeletal genes, including different regulatory factors. These genes showed opposing effects when compared to *bet-1* loss of function animals and were largely dependent on the HAT, *mys-1*. Importantly, the transcriptional changes induced by BET-1 were largely distinct from the UPR^MT^, UPR^ER^, HSR, and the oxidative stress response. However, while we did observe an overlap with a mutant with reduced *daf-2* function, a third of the genes induced by BET-1 were unique and included cytoskeletal and chromatin-related genes, suggesting that at least part of BET-1’s transcriptional program is independent of DAF-2/DAF-16.

In both mammals and *C. elegans*, BET-1/BRD4 has been shown to directly bind chromatin (Dey et al., 2000; Floyd et al., 2013; Shibata et al., 2010) and play an active role in gene expression (Fisher et al., 2013; Lovén et al., 2013). In mammals, BRD4 has direct chromatin decompaction activity. Specifically, BRD4 can act as a HAT to directly acetylate histones H3 and H4, resulting in nucleosome clearance and chromatin decompaction (Devaiah et al., 2016). BRD4 can also recruit transcription super-complexes that promote RNA-PolII activity to stimulate transcription (Donati et al., 2018). While mammalian research has progressed in characterizing a mechanism whereby BRD4 impacts transcription, work on how BET-1 modulates transcription in *C. elegans* has yet to be discovered. Since both BRD4 and BET-1 bind chromatin at histones acetylated at similar lysine residues (H3 K14 or H4 K5/K12 in mammals (Chiang, 2009; Dey et al., 2000) H4 K5, K8, K12, and K16 in *C. elegans* (Shibata et al., 2010)) through conserved bromodomains, it is feasible that the mechanisms of action are similar between mammals and worms. However, additional work is necessary to uncover the direct mechanism whereby BET-1 modulates transcription.

Together, our cross-species approach identified a unique function for BET-1 in actin cytoskeletal maintenance during aging, likely through a conserved function as a transcriptional regulator. Whether BET-1 can directly sense perturbations to cytoskeleton health, or whether the information is relayed by an upstream cytoplasmic sensor, remains to be explored. These findings assign BET-1 a key role as a regulator of cytoskeleton homeostasis, possibly linking previous associations with disease states to this underlying cellular function.

## AUTHOR CONTRIBUTIONS

G.G. and R.H.S. designed all experiments, performed or oversaw all experiments, and prepared the figures and manuscript. R.B.Z. performed computational analysis, figure construction, and writing for all transcriptomics data. C.K.T., E.A.M., and O.S. performed all computational analysis for CRISPR-Cas9 screening. N.D. performed RT-PCR analysis and assisted with lifespans. D.M. and A.A. assisted with motility assays. M.A. and T.C.T. assisted with lifespan assays. M.A.T. performed essential experiments that assisted in development of the manuscript. All authors edited the manuscript.

## DATA AVAILABILITY

All data required to evaluate the conclusions in this manuscript are available within the manuscript and Supplementary Materials. All strains synthesized in this manuscript are derivatives of N2 or other strains from CGC and are either made available on CGC or available upon request. All raw datasets, including CRISPR-Cas9 and RNA-seq, are available through Annotare 2.0 Array Express Accession E-MTAB-11786.

## ACKNOWLEDGEMENTS

We are grateful to Drs. Andrew Dillin and Sean Curran and all members of the Dillin and Curran labs for technical support and sharing of reagents and equipment. GG is supported by T32AG052374, RBZ is supported by the Larry L. Hillblom Foundation Fellowship 019-A-023, C.K.T. is supported by F32AG069388 from the NIA, OS is supported by through the National Institute of General Medical Sciences, and RHS is supported by R00AG065200 from the National Institute on Aging. Some strains were provided by the CGC, which is funded by the NIH Office of Research Infrastructure Programs (P40 OD010440).

## METHODS

### Culturing BJ fibroblasts and cytochalasin screen

The immortalized human foreskin fibroblast line BJ ATCC CRL-2522 (BJ fibroblasts) expressing hTERT and Cas9 were used for the CRISPR-Cas9 based screen as previously described (Schinzel et al., 2019). Cells were cultured in gelatin-coated dishes in DMEM, 15% fetal bovine serum (FBS), 1% glutamax, 1% non-essential amino acids (NEAA), and 1% penicillin/streptomycin. For splitting, cells were washed with PBS, trypsinized, and replated at 1:3 or 1:6 ratios based on confluence. Fresh media was applied every other day.

CellTiter-Glo was used to estimate cell density for titration of cytochalasin D as previously described (Schinzel et al., 2019). Briefly, cells were treated with indicated concentrations of cytochalasin D or a DMSO vehicle control. Plates were washed 1x with PBS to eliminate excess media and CellTiter-Glo media was added to each cell in a 1:3 dilution into cell/PBS mix. Mix was incubated for 30 min at 37°C and luminescence was measured using a Tecan M1000.

For the CRISPR-Cas9 screen, cells were transduced with the AVANA genome-wide sgRNA lentiviral library (Doench et al., 2016; Shalem et al., 2014) and selected for 2 weeks with puromycin to maximize genome editing and target protein depletion. Cells were them split into a control (DMSO) arm and 0.1 µM cytochalasin D treatment arm and harvested after 2 weeks of treatment for sequencing. Genomic DNA (gDNA) extraction was performed on a frozen cell pellet in a 15 mL conical tube on 3×10^7^ – 5×10^7^ cells. Cells were lysed in 50 mM Tris, 50 mM EDTA, 1% SDS, 0.1 mg/mL Proteinase K overnight at 55 °C. Then RNAse A was added to a final concentration of 50 µg/mL and incubated at 37 °C for 30 min. Next, ammonium acetate was added to a final concentration of 2.5 M to precipitate proteins, samples were vortexed at max speed for 20 s and centrifuged at 4,000 x g for 10 min. The supernatant was carefully removed and genomic DNA was extracted with cold 100% isopropanol, washed with cold 70% ethanol, air dried for 30 min to remove excess ethanol, and resuspended in Tris-EDTA. Sequencing was performed using Illumina Next Generation Sequencing as previously described (Shalem et al., 2014). Raw sgRNA counts are provided in **Table S6**.

### C. elegans strains and maintenance

All strains used in this study are derivatives of the N2 wild-type worm from the Caenorhabditis Genetics Center (CGC) and are listed below. Worms are maintained at 15 °C on OP50 *E. coli* B strain, and switched to growth at 20 °C on HT115 *E. coli* K strain for all experimentation. HT115 bacteria carrying a pL4440 empty vector control or expressing double-stranded RNA containing the sequence against a specific target gene were used for all experimentation. Experiments are performed on age-matched animals synchronized using a standard bleaching protocol (Bar-Ziv et al., 2020). Briefly, animals are collected using M9 solution (22 mM KH_2_PO_4_ monobasic, 42.3 mM Na_2_HPO_4_, 85.6mM NaCl, 1 mM MgSO_4_) and bleached in 1.8% sodium hypochlorite and 0.375M KOH diluted in M9 until all carcasses were digested. Intact eggs were then washed 4x with M9 solution followed by L1 synchronization by floating eggs in M9 overnight in a 20 °C incubator on a rotator for a maximum of 16 hours. Synchronized animals are always grown on standard RNAi plates (1 mM CaCl_2_, 5 µg/mL cholesterol, 25 mM KPO_4_, 1 mM MgSO_4_, 2% agar w/v, 0.25% Bacto-Peptone w/v, 51.3 mM NaCl, 1 µM IPTG, and 100 µg/mL carbenicillin; HT115 *E. coli* K strain containing pL4440 vector control or pL4440 with RNAi of interest).

For *bet-1* overexpression, isoforms A, B, and C were defined as per (Shibata et al., 2010) and sequences are provided below. Coding sequences were cloned from cDNA, the endogenous *bet-1* promoter was cloned from gDNA, and an unc-54 3’UTR was cloned from gDNA. Plasmids were injected into N2 worms using a standard microinjection protocol as described (Garcia et al., 2022) with 10 ng/µL of overexpression plasmid, 2.5 ng/µL of pEK2 (*myo-2p::tdtomato*) as a co-injection marker, and 100 ng/µL of pD64 vehicle as filler DNA. Worms positive for the fluorescent pharyx were selected to identify stable arrays. Integration was performed by gamma irradiation where L4 worms were irradiated with 4000-4400 rems of radiation and integration events were selected by finding animals that maintained 100% frequency of co-injection marker in the F3 generation. Lines were then backcrossed into N2 a minimum of 8x to eliminate mutations. For overexpression of *3xHA::GFP::bet-1B*, a GFP sequence containing introns and a 3xHA cassette was cloned upstream of the *bet-1B* coding sequence. Injection and integrations were performed by SUNY Biotech.

To synthesize the *bet-1(uth41)* mutant line, we used a Cas9-RNA protocol as published on the IDT website via the Dernberg lab. Briefly, a mixture of 15 µM trRNA, and 22 µM crRNA (accatgggcaagtccgcgac) were incubated for 5 min at 95 °C, cooled, then mixed with 24 µM Cas9 protein and left at room temperature for 5 °C. 2.5 ng/µL of pEK2 (*myo-2p::tdtomato*) was used added as a co-injection marker and this mixture was injected into *C. elegans* gonads. All progeny positive for the co-injection marker were selected, then sequenced for INDELs that incorporated a premature stop codon. The *bet-1(uth41)* mutant has a premature stop codon at amino acid 17.

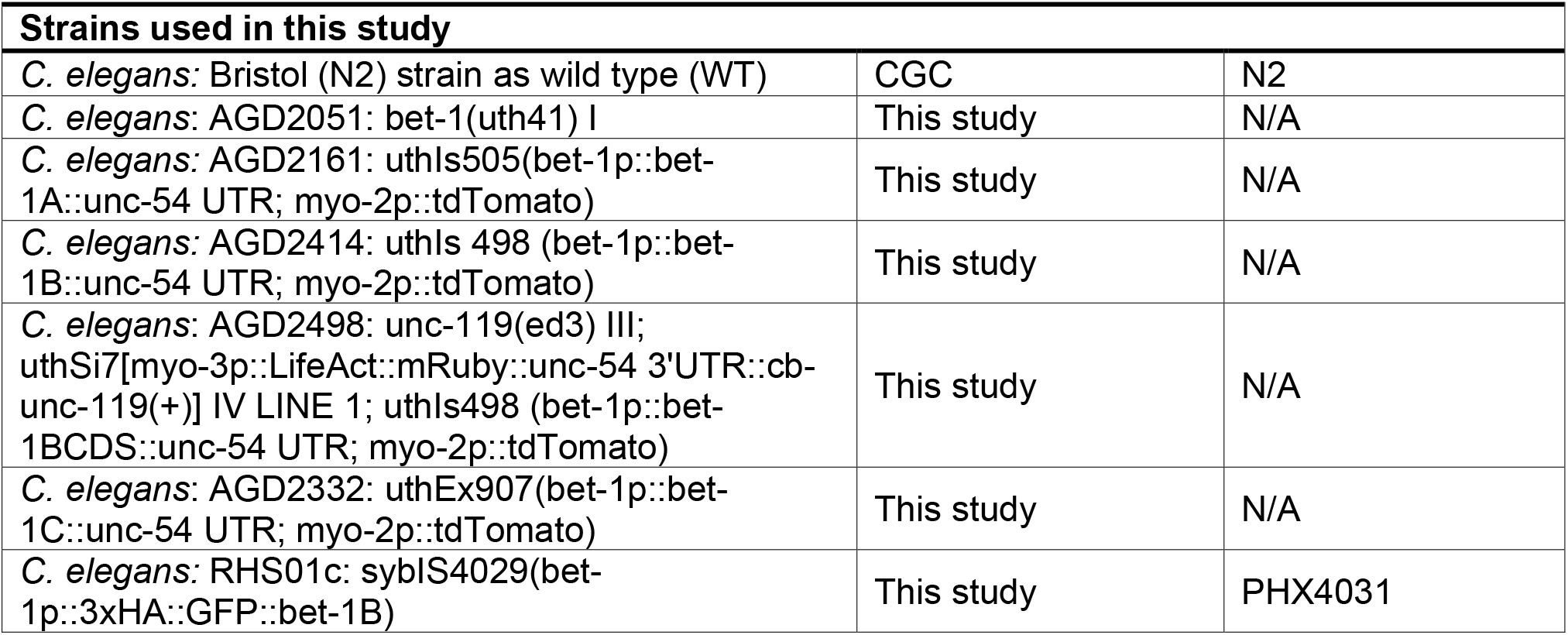

### bet-1A

ATGTCTGAGGGCAGCGGAGACCAATCACAACAACGACCATGGGCAAGTCCGCGACAGCAACCAATCAAAGGAATCGTACAGCCACGAGTACTTCCACCATTCGGAAAGCCAACACGACACACAAACAAACTGGACTACATTATGACAACAGTACTCAAAGAGGCTGGAAAACATAAACATGTCTGGCCGTTTCAGAAGCCCGTCGATGCGGTTGCTTTATGTATTCCTCTATATCACGAGAGAGTCGCCCGACCAATGGACTTGAAAACAATCGAGAATAGACTGAAAAGTACTTATTACACATGTGCTCAAGAATGCATTGATGATATCGAAACAGTTTTCCAAAACTGCTACACATTCAATGGGAAAGAGGACGACGTGACAATTATGGCCCAAAATGTGCACGAAGTGATAAAAAAGTCACTGGAACAAGCACCTCGCGAAGAGCATGATATGGATGTTTATTGGGGAAAAAATAAGAAAAAACCGGCAAAAAGTGACGGTGGATCGAAATCTTCGTCGAGCAAGAAGAATGATGCTCGTGGACCATCTGAAGCACCGTCAGAGGCTGGAAGTGAAGTTTCGTCTGTAACAACAGCATCAGCAGCAGCCCCGACGGTTTCTGAGTCTGCGAGTGTTGCCGCGAAGCCAGAACGAAAAGTGGCCGGAAAGAAGACGGGAAAACGAAAAGCCGAATCAGAAGATGACGAGAAGCCGGAACCTTTGAGAGCAAAACGAGAGGTGGCTGTTGTCAAAAAAGAAGTTCATCAGCCATTGCTCCCAAGTATGAAGCCCTGTCTGAAGCTGCTCAATGATTTTTCTACAAAAAAATATCAGGAATTTGCTTGGCCATTCAACGAACCAGTAGACGCTGAACAACTGGGACTCCATGATTATCATAAAATTATCAAAGAACCAATGGATCTGAAATCAATGAAAGCAAAAATGGAAAGTGGAGCATACAAGGAACCTTCAGATTTCGAGCATGATGTTCGTTTAATGCTCAGGAATTGTTTTCTTTATAATCCAGTCGGTGATCCGGTTCACAGTTTTGGTCTTAGGTTTCAAGAAGTTTTTGATAGACGATGGGCTGAACTAGGTGATTCGAGTTCTCGTGCTTCATCAGTTGCACCTCAATCAGCTCCGATTGCTCCAACTCCGAAAGTAGCAAAATCAAGTGCTCCAAAAGAACCGAAAGAGTCTCGAAAAGAGCATAAAA AGGAGACGACTTTTGAAGCAAGCGGTGCAAAATCGGAGGATTTAATGCAGATAAACAACGCGTTGAGCATGATTCGAGAACGTGAGGAAAAGCTTAAAGCAGAGCTCGCCGCTGCACAAGC GATAAAGGATAAACTGACGAGTGTGAAGAATCGACGAGAAGATAATCCGAATGAGCCATTTCCGGAGAAGCTTATCAATGAGACAAGAGCCTTGTGCACGACGCAAGTTGGACAAAATGCTTCAAGTTCTTCAGCTTCTTCTGCTGCTTTGAGGAACGGACGAAGCAAAAAAGCAGCATCCGCACGTCTCTATGGTTACGAATTTGATTCGGATGATGAGGATAATAAGATGGCACTGACTTATGAGGAAAAACGAAACTTGAGCAATCTGATTAATAATTTACCCAACAATCAACTCAACACCATAATTTCGATTATTCAACGGAGAGAACGAAGCGCTCTGATGCAACAACAACTCGATGACAGTGAGGTTGAACTGGATTTCGAATCACTTGGAGATATGTGCCTGAGAGAAATGGGTGCATTTATCAAAACAATTCCAACATTAAACGGAAATGGCGATGATGAGAAGCCGAAAACGTCTTCGAATCCGACATCTTCTGGAGCAACAGGATCAAAGGGTTCGTCGTCGTTGGAGAGCAAAAATGGAAAGAAAAAGAAAAACTTCAATATGTCCGAATCCTCGGATGATGAGACGTCGAATAGTCGAAAACGTCGAAAGAGAGAGAGCAGTGAATCACAGAGCTCTTCGTCCAGTGATGATGATTCAGATGATGAGGATAGGCCGAGTATTCCCCGTAAATCAGGTCAACCACCATCAACATCACGTGAATGGAATCAATCATCAGCTCCTCCACCACGAATGGGAGGAATGGGAGGACAACCACCAATGTCACGAGTACCTGCATCATCATCCACATCTGTATCAGCAATCGGAAAGAACAACGCAGCCGCCTCGTCGAATTCATATCAAGCTCCAAAACCTGCACCAGTACCAGCACCAACATCATCAAGACCTCCGGCAGCACCGAGACCACCGTCAAAACCAAAGAAAACGGGTGGAGCGAGTATTCTTGATACTCTACTTCCAGATACATTTGGAGCATCACCTCCCCAGTTTTTCCAGTCGCAACCAACAACGTCGGCTACGATTAGATCACCAACGGAAAGCCAACCCGGGAATGGTGAAGACGAGCAGACCAGGATTCAGAGGATGCGGATGGAGGCAAAGCGAGCCCGCCAAAAAGAAGACGAAGGCAGTGTCTCGTTGTCAAACCAAATGGAAATGATGGCTGCATTTGAATTTGATAATACATA TTAA

### bet-1B

ATGTCTGAGGGCAGCGGAGACCAATCACAACAACGACCATGGGCAAGTCCGCGACAGCAACCAATCAAAGGAATCGTACAGCCACGAGTACTTCCACCATTCGGAAAGCCAACACGACACACAAACAAACTGGACTACATTATGACAACAGTACTCAAAGAGGCTGGAAAACATAAACATGTCTGGCCGTTTCAGAAGCCCGTCGATGCGGTTGCTTTATGTATTCCTCTATATCACGAGAGAGTCGCCCGACCAATGGACTTGAAAACAATCGAGAATAGACTGAAAAGTACTTATTACACATGTGCTCAAGAATGCATTGATGATATCGAAACAGTTTTCCAAAACTGCTACACATTCAATGGGAAAGAGGACGACGTGACAATTATGGCCCAAAATGTGCACGAAGTGATAAAAAAGTCACTGGAACAAGCACCTCGCGAAGAGCATGATATGGATGTTTATTGGGGAAAAAATAAGAAAAAACCGGCAAAAAGTGACGGTGGATCGAAATCTTCGTCGAGCAAGAAGAATGATGCTCGTGGACCATCTGAAGCACCGTCAGAGGCTGGAAGTGAAGTTTCGTCTGTAACAACAGCATCAGCAGCAGCCCCGACGGTTTCTGAGTCTGCGAGTGTTGCCGCGAAGCCAGAACGAAAAGTGGCCGGAAAGAAGACGGGAAAACGAAAAGCCGAATCAGAAGATGACGAGAAGCCGGAACCTTTGAGAGCAAAACGAGAGGTGGCTGTTGTCAAAAAAGAAGTTCATCAGCCATTGCTCCCAAGTATGAAGCCCTGTCTGAAGCTGCTCAATGATTTTTCTACAAAAAAATATCAGGAATTTGCTTGGCCATTCAACGAACCAGTAGACGCTGAACAACTGGGACTCCATGATTATCATAAAATTATCAAAGAACCAATGGATCTGAAATCAATGAAAGCAAAAATGGAAAGTGGAGCATACAAGGAACCTTCAGATTTCGAGCATGATGTTCGTTTAATGCTCAGGAATTGTTTTCTTTATAATCCAGTCGGTGATCCGGTTCACAGTTTTGGTCTTAGGTTTCAAGAAGTTTTTGATAGACGATGGGCTGAACTAGGTGATTCGAGTTCTCGTGCTTCATCAGTTGCACCTCAATCAGCTCCGATTGCTCCAACTCCGAAAGTAGCAAAATCAAGTGCTCCAAAAGAACCGAAAGAGTCTCGAAAAGAGCATAAAAAGGAGACGACTTTTGAAGCAAGCGGTGCAAAATCGGAGGATTTAATGCAGATAAACAACGCGTTGAGCATGATTCGAGAACGTGAGGAAAAGCTTAAAGCAGAGCTCGCCGCTGCACAAGC GATAAAGGATAAACTGACGAGTGTGAAGAATCGACGAGAAGATAATCCGAATGAGCCATTTCCGGAGAAGCTTATCAATGAGACAAGAGCCTTGTGCACGACGCAAGTTGGACAAAATGCTT CAAGTTCTTCAGCTTCTTCTGCTGCTTTGAGGAACGGACGAAGCAAAAAAGCAGCATCCGCACGTCTCTATGGTTACGAATTTGATTCGGATGATGAGGATAATAAGATGGCACTGACTTATGAGGAAAAACGAAACTTGAGCAATCTGATTAATAATTTACCCAACAATCAACTCAACACCATAATTTCGATTATTCAACGGAGAGAACGAAGCGCTCTGATGCAACAACAACTCGATGACAGTGAGGTTGAACTGGATTTCGAATCACTTGGAGATATGTGCCTGAGAGAAATGGGTGCATTTATCAAAACAATTCCAACATTAAACGGAAATGGCGATGATGAGAAGCCGAAAACGTCTTCGAATCCGACATCTTCTGGAGCAACAGGATCAAAGGGTTCGTCGTCGTTGGAGAGCAAAAATGGAAAGAAAAAGAAAAACTTCAATATGTCCGAATCCTCGGATGATGAGACGTCGAATAGTCGAAAACGTCGAAAGAGAGAGAGCAGTGAATCACAGAGCTCTTCGTCCAGTGATGATGATTCAGATGATGAGGATAGGCCGAGTATTCCCCGTAAATCAGGTCAACCACCATCAACATCACGTGAATGGAATCAATCATCAGCTCCTCCACCACGAATGGGAGGAATGGGAGGACAACCACCAATGTCACGAGTACCTGCATCATCATCCACATCTGTATCAGCAATCGGAAAGAACAACGCAGCCGCCTCGTCGAATTCATATCAAAAATTTTATAATTGTTTTCACAGTTATACTCCACCTTTAAAAGTTGAAAAAAAAATCATCAAATTACTGGTAAATTTTTGTTAA

### bet-1C

ATGTCTGAGGGCAGCGGAGACCAATCACAACAACGACCATGGGCAAGTCCGCGACAGCAACCAATCAAAGGAATCGTACAGCCACGAGTACTTCCACCATTCGGAAAGCCAACACGACACACAAACAAACTGGACTACATTATGACAACAGTACTCAAAGAGGCTGGAAAACATAAACATGTCTGGCCGTTTCAGAAGCCCGTCGATGCGGTTGCTTTATGTATTCCTCTATATCACGAGAGAGTCGCCCGACCAATGGACTTGAAAACAATCGAGAATAGACTGAAAAGTACTTATTACACATGTGCTCAAGAATGCATTGATGATATCGAAACAGTTTTCCAAAACTGCTACACATTCAATGGGAAAGAGGACGACGTGACAATTATGGCCCAAAATGTGCACGAAGTGATAAAAAAGTCACTGGAACAAGCACCTCGCGAAGAGCATGATATGGATGTTTATTGGGGAAAAAATAAGAAAAAACCGGCAAAAAGTGACGGTGGATCGAAATCTTCGTCGAGCAAGAAGAATGATGCTCGTGGACCATCTGAAGCACCGTCAGAGGCTGGAAGTGAAGTTTCGTCTGTAACAACAGCATCAGCAGCAGCCCCGACGGTTTCTGAGTCTGCGAGTGTTGCCGCGAAGCCAGAACGAAAAGTGGCCGGAAAGAAGACGGGAAAACGAAAAGCCGAATCAGAAGATGACGAGAAGCCGGAACCTTTGAGAGCAAAACGAGAGGTGGCTGTTGTCAAAAAAGAAGTTCATCAGCCATTGCTCCCAAGTATGAAGCCCTGTCTGAAGCTGCTCAATGATTTTTCTACAAAAAAATATCAGGAATTTGCTTGGCCATTCAACGAACCAGTAGACGCTGAACAACTGGGACTCCATGATTATCATAAAATTATCAAAGAACCAATGGATCTGAAATCAATGAAAGCAAAAATGGAAAGTGGAGCATACAAGGAACCTTCAGATTTCGAGCATGATGTTCGTTTAATGCTCAGGAATTGTTTTCTTTATAATCCAGTCGGTGATCCGGTTCACAGTTTTGGTCTTAGGTTTCAAGAAGTTTTTGATAGACGATGGGCTGAACTAGGTGATTCGAGTTCTCGTGCTTCATCAGTTGCACCTCAATCAGCTCCGATTGCTCCAACTCCGAAAGTAGCAAAATCAAGTGCTCCAAAAGAACCGAAAGAGTCTCGAAAAGAGCATAAAAAGGAGACGACTTTTGAAGCAAGCGGTGCAAAATCGGAGGATTTAATGCAGATAAACAACGCGTTGAGCATGATTCGAGAACGTGAGGAAAAGCTTAAAGCAGAGCTCGCCGCTGCACAAGCGATAAAGGATAAACTGACGAGTGTGAAGAATCGACGAGAAGATAATCCGAATGAGCCATTTCCGGAGAAGCTTATCAATGAGACAAGAGCCTTGTGCACGACGCAAGTTGGACAAAATGCTTCAAGTTCTTCAGCTTCTTCTGCTGCTTTGAGGAACGGACGAAGCAAAAAAGCAGCATCCGCACGTCTCTATGGTTACGAATTTGATTCGGATGATGAGGATAATAAGATGGCACTGACTTATGAGGAAAAACGAAACTTGAGCAATCTGATTAATAATTTACCCAACAATCAACTCAACACCATAATTTCGATTATTCAACGGAGAGAACGAAGCGCTCTGATGCAACAACAACTCGATGACAGTGAGGTTGAACTGGATTTCGAATCACTTGGAGATATGTGCCTGAGAGAAATGGGTGCATTTAT CAAAACAATTCCAACATTAAACGGAAATGGCGATGATGAGAAGCCGAAAACGTCTTCGAATCCGACATCTTCTGGAGCAACAGGATCAAAGGGTTCGTCGTCGTTGGAGAGCAAAAATGGA AAGAAAATAA

### bet-1p

CACAGGTCTCTAGTGTATCCACTTCGAATGCGATGCCCGAAACCTCTTCATCCATCCGTCTCCTTCTCGCTCTCTCTCTCTCTCTCTCTTCTCCATCTCTCTCCACATTTTGCCTGCTATCTCGTGATTGTCGTCCCGTCCGTGTTCCGCCGCACACACTGCCTGTCTTCTCTTAACCGTGTGTCGATCAACTCCCAAACCGCTACGCTATTTCTCTCTCCCTCTCTCTCTCTCTTCGGCGGTGACATTTCTGACTAGATGGTCATACAAAACGCGTGCTGCGCGCGCGCTCCGCAAAAATCGACGCGAATCGATTAATGTGCGTCTCGTTTCTCTATCTCTGACCGCCCCCGCTTCAACCTAACACTATTTTTGAATGCTTTTCAACTGTAACTTGCAGCTAATTAGAAGTTGAGAGATAACCTGTTGCGATTGGCTCCGGGCAAGGGTTGGGAGGTCGCACCAGAAATTTTAGAGCTCTAGGATTTCAAATTTTTGGGTTTCAAGACCGTAACATGATTTTCTTGGAAATTTATCACAAATCATGTAGAAAATCGATATCAGTAAGAGGGAGTGAGTGATCTATCATTTTTTATCTTTCGATCTGAAATTCCACAGCGAAGGTTTTCTGCCGAAATTTCGAAATTGGTATTTTGAACTATCCGATAATTCGTAGAACATCAAGATAAAGTGTCAACCTATAGAAAATCACATGATTCGTCAGAAAATAACATTAATTTCATATGAAATAGTTGAGAAAGTGCTCAAAAATGGCCTAAAATTATCCAATAATCGACATTTGACAACTTTCAGCACACTTTTGAACCGTTTATCAATTGTTTCTGCTGAAATAGACGTATTTTTCGGACGAATCGAGTGATTTCCTATAGTTTTACACTGATTTTTGACAAAAAAATATTGATAGAACATGGTGCATTAGGCAATTTTTTAGAATTGCCGTCTACACCTGATTTCGATGGGTCCTCGTGACAAGACCCAAAATTTTATTATTTTTATCGTTGAAAAAAATCAAATCAATAACACCGCAATCACCATTTGCAAAGTTTAATTAAATACAATTTTTATTAAAATATTTCAGGAATAAAAATATTAGTCAGAATAATCCCATGTTCTTCTAGCGATTTCAACTAATTCTTTGAAAATAACATTTCTTGGAATTTAAGAATACGAAAATAGTCACTTCTTTGTATTCTAGAAACGCTAATTCCTGCAACCGACAAATTAAAAGTACAAAAAATGATACGGCAAGCGCGCTCCAATTCAAATCGAGTCTCCCGCCTTCCTTGACGTCATTGCTAACAGCTGCTTCGGTTTTTTCCTCCAAATTTCGTGGTTCAAATTTTATTTTTAATTGAATTTTAACAAAATAGGAAGCTAGTTGAGTAACATTTATTATTAATTTTGTAAAATATTCTGCAAATTCGGCGTTTTCTTTTAATTCAAATAAAAGTTTTCAATAAAAAAAATCGATAT TTTCAG

### unc-54 3’ UTR

CATCTCGCGCCCGTGCCTCTGACTTCTAAGTCCAATTACTCTTCAACATCCCTACATGCTCTTTCTCCCTGTGCTCCCACCCCCTATTTTTGTTATTATCAAAAAACTTCTCTTAATTTCTTTGTTTTTTAGCTTCTTTTAAGTCACCTCTAACAATGAAATTGTGTAGATTCAAAAATAGAATTAATTCGTAATAAAAAGTCGAAAAAAATTGTGCTCCCTCCCCCCATTAATAATAATTCTATCCCAAAATCTACACAATGTTCTGTGTACACTTCTTATGTTTTTTACTTCTGATAAATTTTTTTGAAACATCATAGAAAAAACCGCACACAAAATACCTTATCATATGTTACGTTTCAGTTTATGACCGCAATTTTTATTTCTTCGCACGTCTGGGCCTCTCATGACGTCAAATCATGCTCATCGTGAAAAAGTTTTGGAGTATTTTTGGAATTTTTCAATCAAGTGAAAGTTTATGAAATTAATTTTCCTGCTTTTGCTTTTTGGGGTTTCCCCTATTGTTTGTCAAGATTTCGAGGACGGCGTTTTTCTTGCTAAAATCACAAGTATTGATGAGCACGATGCAAGAAAGATCGGAAGAAGGTTTGGGTTTGAGGCTCAGTGGAAG

### GFP (introns are lower case, stop codon removed)

ATGAGTAAAGGAGAAGAACTTTTCACTGGAGTTGTCCCAATTCTTGTTGAATTAGATGGTGA TGTTAATGGGCACAAATTTTCTGTCAGTGGAGAGGGTGAAGGTGATGCAACATACGGAAAACTTACCCTTAAATTTATTTGCACTACTGGAAAACTACCTGTTCCATGGgtaagtttaaacatatatatactaactaaccctgattatttaaattttcagCCAACACTTGTCACTACTTTCTGTTATGGTGTTCAATGCTTCTC GAGATACCCAGATCATATGAAACGGCATGACTTTTTCAAGAGTGCCATGCCCGAAGGTTAT GTACAGGAAAGAACTATATTTTTCAAAGATGACGGGAACTACAAGACACgtaagtttaaacagttcggtactaactaaccatacatatttaaattttcagGTGCTGAAGTCAAGTTTGAAGGTGATACCCTTGTTAATAGA ATCGAGTTAAAAGGTATTGATTTTAAAGAAGATGGAAACATTCTTGGACACAAATTGGAATA CAACTATAACTCACACAATGTATACATCATGGCAGACAAACAAAAGAATGGAATCAAAGTTgtaagtttaaacatgattttactaactaactaatctgatttaaattttcagAACTTCAAAATTAGACACAACATTGAAGATG GAAGCGTTCAACTAGCAGACCATTATCAACAAAATACTCCAATTGGCGATGGCCCTGTCCTTTTACCAGACAACCATTACCTGTCCACACAATCTGCCCTTTCGAAAGATCCCAACGAAAAGAGAGACCACATGGTCCTTCTTGAGTTTGTAACAGCTGCTGGGATTACACATGGCATGGATGAACTATACAAA

### 3xHA

TATCCATATGACGTGCCGGACTACGCGTACCCGTATGATGTTCCAGACTACGCCTATCCGTACGACGTACCAGATTATGCA

### bet-1 RNAi

CAGCAACCAATCAAAGGAATCGTACAGCCACGAGTACTTCCACCATTCGGAAAGCCAACACGACACACAAACAAACTGGACTACATTATGACAACAGTACTCAAAGAGGCTGGAAAACATAAACATGTCTGGCCGTTTCAGAAGCCCGTCGATGCGGTTGCTTTATGTATTCCTCTATATCACGAGAGAGTCGCCCGACCAATGGACTTGAAAACAATCGAGAATAGACTGAAAAGTACTTATTACACATGCGCTCAAGAATGCATTGATGATATCGAAACAGTTTTCCAAAACTGCTACACATTCAATGGGAAAGAGGACGACGTGACAATTATGGCCCAAAATGTGCACGAAGTGATAAAAAAGTCACTGGAACAAGCACCTCGCGAAGAGCATGATATGGATGTTTATTGGGGAAAAAATAAGAAAAAACCGGCAAAAAGTGACGGTGGATCGAAATCTTCGTCGAGCAAGAAGAATGATGCTCGTGGACCATCTGAAGCACCGTCAGAGGCTGGAAGTGAAGTTTCGTCTGTAACAACAGCATCAGCAGCAGCCCCGACGGTTTCTGAGTCTGCGAGTGTTGCCG

### mys-1 RNAi

AAAAAAGCAGGCTTGACCGAGCCGAAGAAGGAGATTATAGAGGACGAAAATCATGGAATATCCAAGAAAATACCAACAGATCCCAGGCAATACGAGAAAGTTACAGAGGGATGCCGGTTATTGGTCATGATGGCTTCACAAGAAGAAGAAAGATGGGCCGAAGTTATTTCAAGATGCCGAGCTGCAAATGGTTCAATTAAATTCTATGTCCATTATATCGATTGCAACCGAAGACTTGACGAATGGGTTCAGTCTGATAGGCTCAATTTAGCGTCGTGTGAGCTACCAAAAAAAGGAGGAAAGAAAGGAGCACACTTGCGGGAAGAAAATCGAGATTCGAATGAAAATGAAGGAAAGAAAAGCGGCCGAAAACGAAAGATTCCACTACTTCCGATGGATGATCTCAAGGCGGAATCCGTAGATCCATTACAAGCAATTTCAACGATGACCAGCGGATCTACTCCAAGTCTTCGAGGTTCCATGTCGATGGTCGGCCATAGTGAAGATGCAATGACAAGGATCCGAAATGTCGAATGCATTGAACTAGGAAGATCACGAATTCAGCCATGGTACTTTGCACCTTATCCACAACAATTGACAAGTTTGGATTGTATTTATATTTGCGAATTTTGTCTGAAATATCTAAAGTCGAAAACTTGTCTGAAACGGCACNTGGAAAAATGTGCAATGTGTCACCCACCTGGCAATCAAATCTACAGTCACGATAAACTTTCATTTTTTGAAATCGACGGCCGCAAAAACAAAAGCTATGCTCAGAATCTATGCCTGCTTGCCA AACTT

### C. elegans screen

*C. elegans* orthologs of human genes from the cytochalasin screen were identified using Ortholist 2 (Kim et al., 2018). RNAis were isolated from the Vidal (Reboul et al., 2001) or Ahringer (Lee et al., 2003) RNA libraries and sequence-verified using standard sanger sequencing. RNAi constructs that matched the expected sequences at a bp length > 150 were included in the screen. For synthetic lethality screens, RNAi cultures were grown in a deep-well 96-well plate to saturation and were mixed at a 10%/90% ratio of *act-1* RNAi to candidate gene RNAi. Animals were grown on these 10%/90% mix and grown at 20 °C and screened at day 1 of adulthood. All hits are defined as those that show any observable difference between the 10%/90% gene/*act-1* mix in comparison to 100% candidate gene RNAi alone. All screening was performed with the researcher blind to the identity of each RNAi and was screened by two independent researchers. Only hits that had phenotypes scored as positive by both researchers were included as hits and images are made available in Fig. 1.

### C. elegans microscopy

Animals were always imaged at the specified ages in figure legends using standard bleaching protocols for synchronization. For all aging experiments, animals were aged on RNAi plates supplemented with FUDR from day 1 of adulthood until the desired stage. 100 µL of 10 mg/mL FUDR were spotted on the bacterial lawn. At least 1 replicate was performed without FUDR and manual picking of animals away from their progeny as previously described (Higuchi-Sanabria et al., 2018) to ensure that measurable effects were independent of FUDR. For all microscopy, representative images of three independent biological replicates are shown.

For live-cell imaging, animals are picked off of plates and mounted directly onto a microscope slide containing M9 + 0.1 M sodium azide. For standard wide-field microscopy, images were acquired on either a Zeiss AxioObserver.Z1 microscope equipped with a lumencor sola light engine and Ziess axiocam 506 camera driven by Zeiss ZenBlue software using a 63x/1.4 PlanApochromat objective, standard dSRed filter (Zeiss filter set 43), and a DFC9000 camera; or a Leica Thunder Imager equipped with a 63x/1.4 Plan AproChromat objective, standard dsRed filter (11525309), Leica DFC9000 GT camera, a Leica LED5 light source, and run on LAS X software.

For imaging of GFP::BET-1, animals were bleached to isolate eggs. 100 µL of egg/M9 mix was mixed with 500 µL of 4% PBS diluted in PBS and fixed at room temperature on a belly dancer for 11 min. Samples were frozen at – 80°C until imaging. Prior to imaging, PFA was washed using 1 mL of PBS and shaking on a belly dancer for 10 min at room temperature. A total of 3x PBS washes was performed. For staining, samples were submerged in 1 mL of PBS, then 0.75 µL of 5 mg/mL DAPI dissolved in DMSO was added. Eggs were incubated with DAPI for 50 min at room temperature on a belly dancer. Excess DAPI was washed with 3x PBS washes at 10 min each at room temperature on the belly dancer. 5 µL of egg/PBS mix was mounted onto a slide and mixed with 5 µL of VectaShield mounting media and imaged on a Stellaris 5 confocal microscope equipped with a white light laser source and 405 nm laser source, HyD detector, 63x/1.4 Plan ApoChromat objective, and run on LAS X software.

### C. elegans RT-PCR and RNA-seq analysis

For RNA isolation, all RNa collection was performed at day 1 of adulthood. ∼1000 animals were harvested from RNAi plates using M9. Animals were pelleted by gravity by allowing adult worms to settle to the bottom of the tube and aspirating off eggs and L1. Animals were washed and gravity settled 3x to remove a majority of progeny, then animals were placed into Trizol solution and worms were freeze/thawed 3x with liquid nitrogen with a 30 sec vortexing step between each freeze cycle. After the final thaw, chloroform was added at a 1:5 chlorform/trizol ratio and aqueous separation of RNA was performed via centrifugation in a heavy gel phase-lock tube (VWR, 10847-802). The aqueous phase was mixed 1:1 with isopropanol then applied to a Qiagen RNeasy Mini Kit (74106) and RNA purification was performed as per manufacturer’s directions.

Library preparation was performed using a Kapa Biosystems mRNA Hyper Prep Kit. Sequencing was performed at the Vincent J Coates Genomic Sequencing Core at the University of California, Berkeley using an Illumina HS4000 mode SR100. Four biological replicates were measured per condition. Reads per gene were quantified using kallisto (Bray et al., 2016), with WBcel235 as the worm reference genome. Fold changes were determined using DESeq2 (Love et al., 2014). XBP-1 gene targets were defined as previously experimentally determined (Urano et al., 2002). GO enrichment was calculated using WormEnrichr (Chen et al., 2013). *bet-1B* overexpression, *bet-1* RNAi, and *bet-1(uth41)* mutants were compared to N2 wildtype control. In addition, *bet-1* overexpressing worms were grown on *mys-1* RNAi and compared to a *mys-1* RNAi control.

For RT-PCR, cDNA synthesis was performed using the QuantaBio cDNA supermix Qscript (101414-102) using 1 µg of RNA. RT-PCR was performed using NEB Q5 DNA polymerase as per manufacturer’s guidelines using primers listed below. Four biological replicates were performed per condition. Image quantification was performed using ImageJ by drawing an ROI of equal size around each band and quantifying for integrated density. Data was normalized to a *tba-1* loading control.

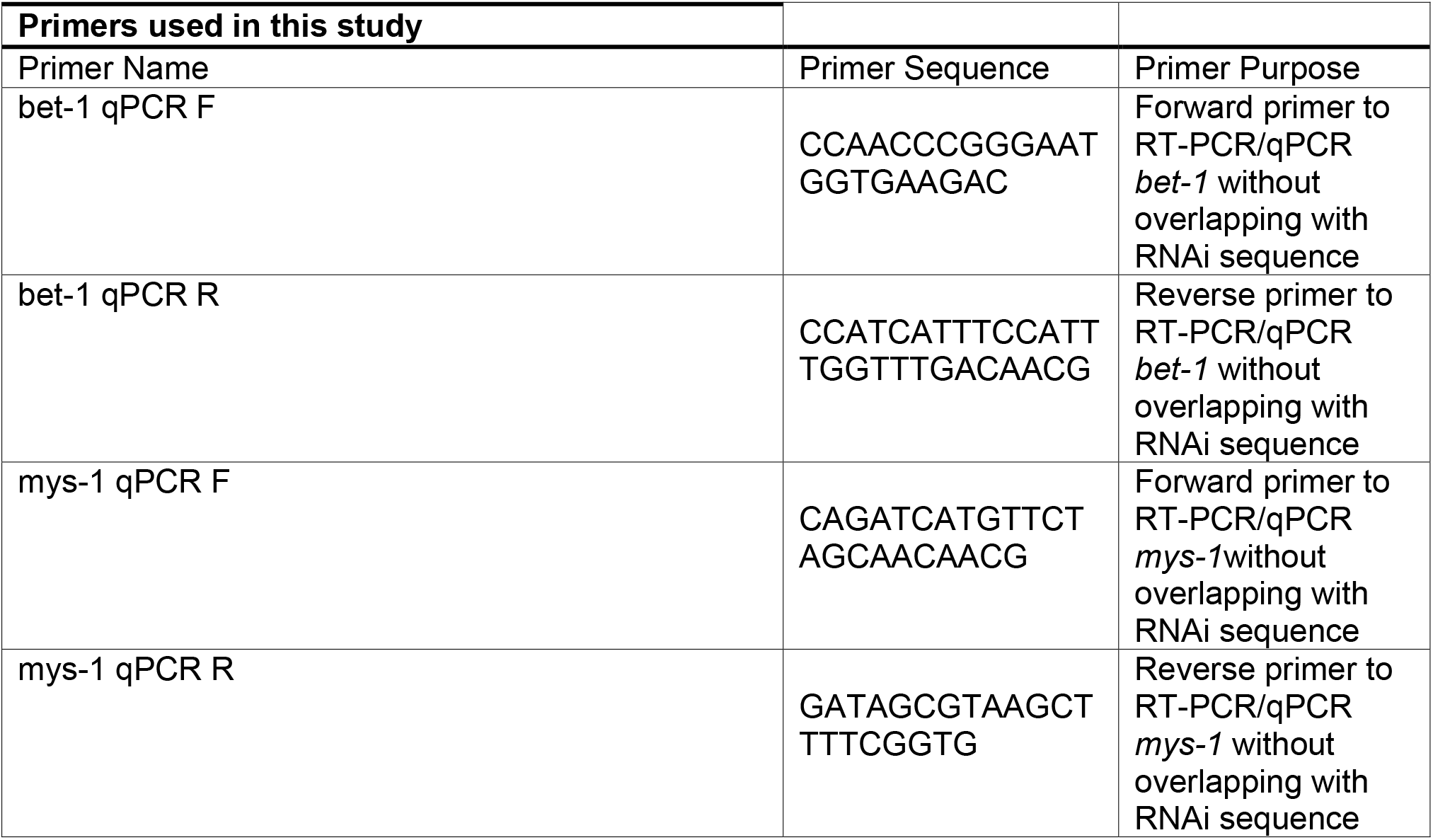
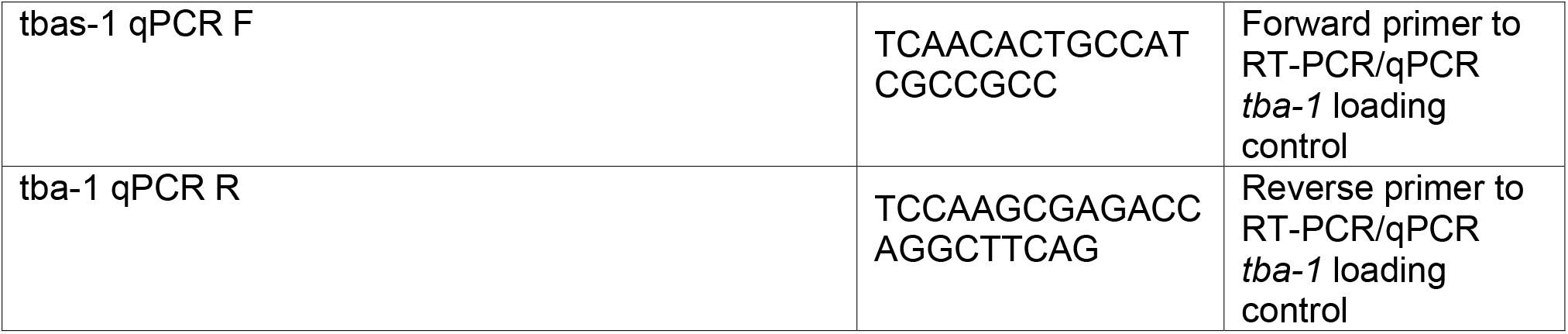

### C. elegans thrashing assay

Thrashing assays were performed on animals synchronized via bleaching and aged on plates containing FUDR from day 1. 100 µL of 10 mg/mL FUDR were spotted on the bacterial lawn. At the desired age, plates containing adult animals were flooded with 100 µL of M9 solution, and 30 sec videos were acquired on an M205FCA stereomicroscope equipped with a Leica K5 microscope and run on LAS X software. Thrashing was measured by eye over a 10 second period. A single trash is defined as bending of >50% of the animal’s body in the opposite direction. Representative data of three independent biological replicates are presented. Dot plots were generated using Prism 7 software where every dot represents a single animal and lines represent median and interquartile range. All statistics were performed using non-parametric Mann-Whitney testing.

### C. elegans brood size assay

A synchronized population of animals were collected via bleaching and 10 L4 animals were moved onto individual plates. Every 12 hours, animals were moved onto fresh plates and plates containing eggs were stored in a 15 ° incubator for 2-3 days. All live progeny on every egg-lay plate were scored and summed to determine brood size. Dot plots were generated using Prism 7 software where every dot represents a single animal and lines represent median and interquartile range. All statistics were performed using non-parametric Mann-Whitney testing.

### C. elegans lifespan assay

*C. elegans* lifespan assays were performed on standard RNAi plates and were all performed at 20 °C as previously described (Bar-Ziv et al., 2020). Adult worms were moved away from progeny daily onto fresh RNAi plates until no progeny were visible (∼7-8 days). Animals were then scored every other day until all animals were scored as either dead or censored. All animals exhibiting bagging, intestinal leaking out of the vulva, or other age-unrelated death were censored and removed from quantification. For lifespans on FUDR, animals were grown on RNAi plates supplemented with FUDR from day 1 of adulthood until the desired stage. 100 µL of 10 mg/mL FUDR were spotted on the bacterial lawn. At least 1 replicate for every lifespan was performed in the absence of FUDR. All statistical analysis was performed using Prism7 software using LogRank testing. All lifespan experiments were performed with researchers blinded to sample conditions. Representative data are represented in figures and all replicates are made available in **Table S3-4**.

## SUPPLEMENTARY FIGURES, TABLES, LEGENDS

**Fig. S1.**
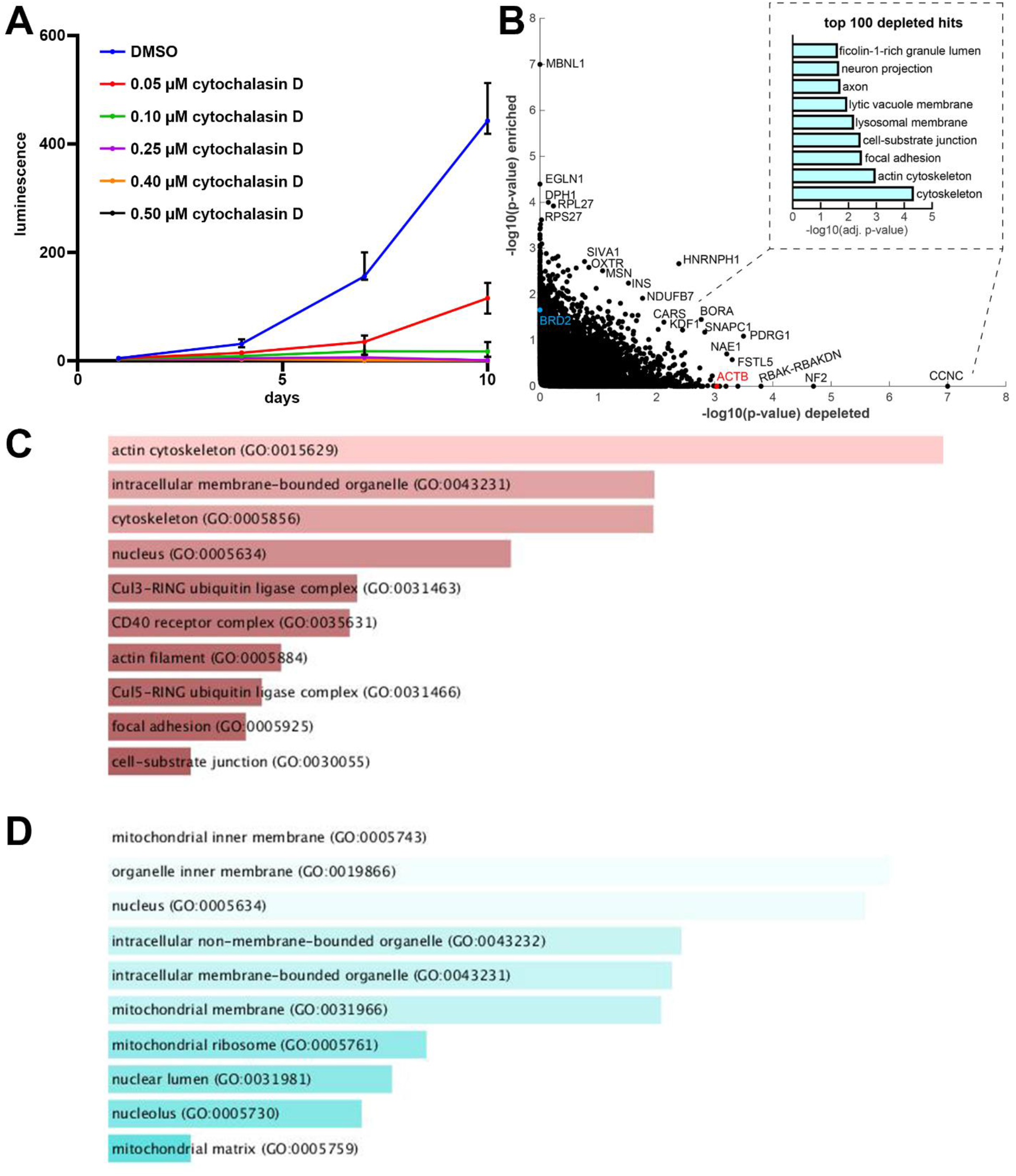
Cytochalasin CRISPR-Cas9 screening. **(A)** Cell density was determined via CellTiter-Glo analysis at day 1, day 4, and day 10 of exposure to indicated concentration of cytochalasin D or DMSO control as described in STAR Methods. n = 4 per condition and data is represented as median +/- interquartile range. **(B)** The p-values (max p-val) of each gene when calculated for enriched and depleted genes, upon Cytochalasin D treatment, is shown. (inset) enrichment analysis for cellular components of the top 100 depleted genes (Chen et al., 2013). All significantly depleted **(C)** and enriched **(D)** genes were defined as those with a p-value < 0.05 and can be found in **Table S1-2**. Gene lists were run through Enrichr and gene ontologies (GO) for cellular component were graphed and sorted by p-value ranking. All raw counts are available on **Table S6**.

**Fig. S2.**
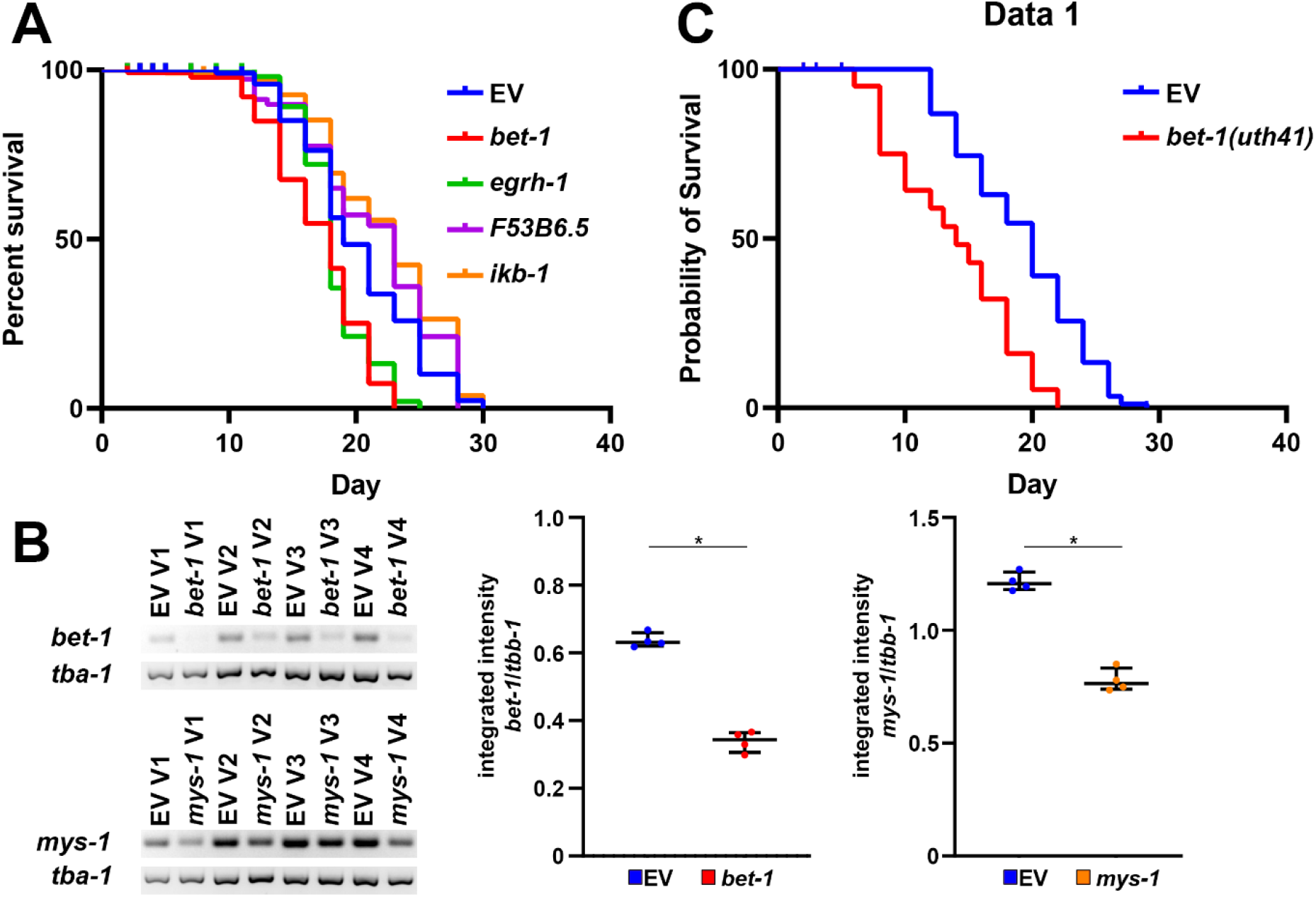
Knockdown and knockout of *bet-1* decrease lifespan. **(A)** Lifespans of wild-type, N2 animals grown on empty vector (EV, blue), *bet-1* (red), *egrh-1* (green), *F53B6.5* (purple), or *ikb-1* (orange) RNAi from hatch. See **Table S3-4** for lifespan statistics. **(B)** RT-PCR of transcripts in N2 animals grown on empty vector (EV), *bet-1*, or *mys-1* RNAi from hatch. RNA was isolated in day 1 adults, followed by cDNA synthesis and PCR. Quantification was performed in ImageJ measuring the integrated intensity of bands and normalizing against a *tba-1* loading control. Left side shows PCR band for each of four biological replicates (V1-V4). Right side are dot plots where each dot represents a single biological replicate and lines represent median and interquartile range. * = p < 0.05 calculated using non-parametric Mann-Whitney testing. **(C)** Lifespans of N2 (EV, blue) and *bet-1(uth41)* (red) animals grown on empty vector RNAi from hatch. See **Table S3-4** for lifespan statistics.

**Fig. S3.**
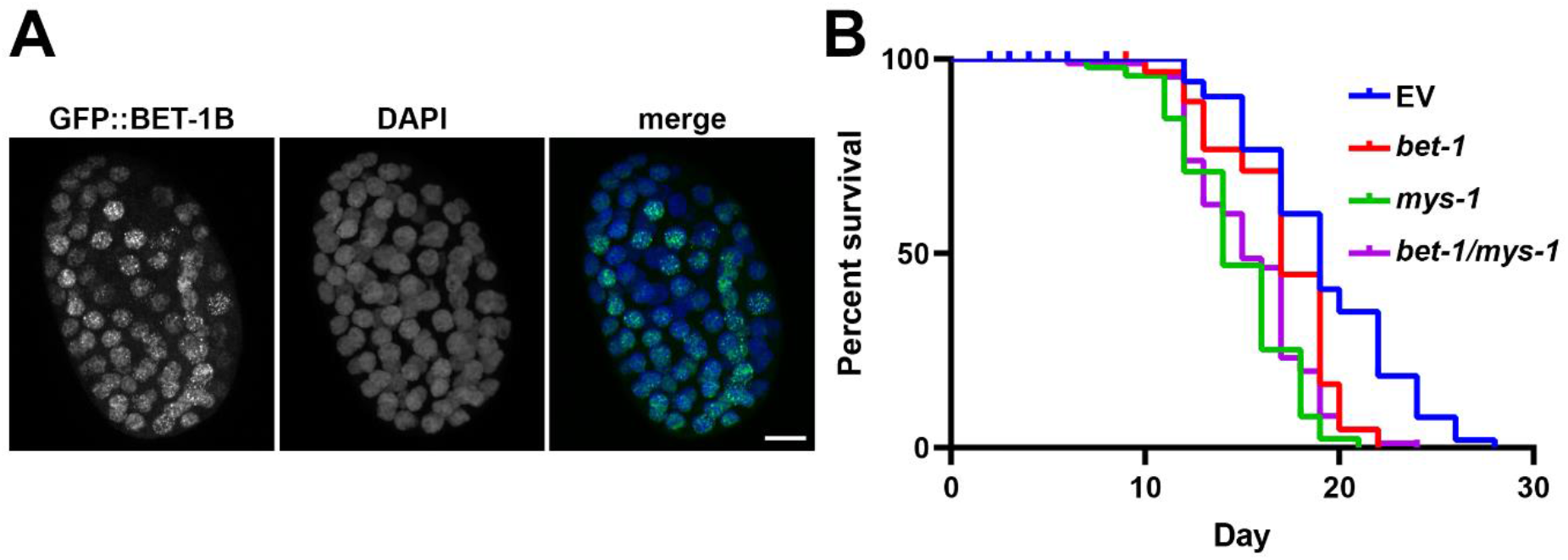
BET-1 functions as a transcriptional regulator. **(A)** Animals overexpressing *GFP::bet-1B* were grown on empty vector RNAi from hatch. Eggs were isolated by a standard bleaching protocol, fixed using 4% PFA, and counterstained with DAPI as described in STAR Methods. Images were collected on a Leica Stellaris 5 confocal. Scale bar is 5 µ. **(B)** Lifespans of wild-type, N2 animals grown on empty vector (EV, blue) or a 50/50 mix of EV/*bet-1* (red), EV/*mys-1* (green), or *bet-1/mys-1* (purple) from hatch. See **Table S3-4** for lifespan statistics.

**Fig. S4.**
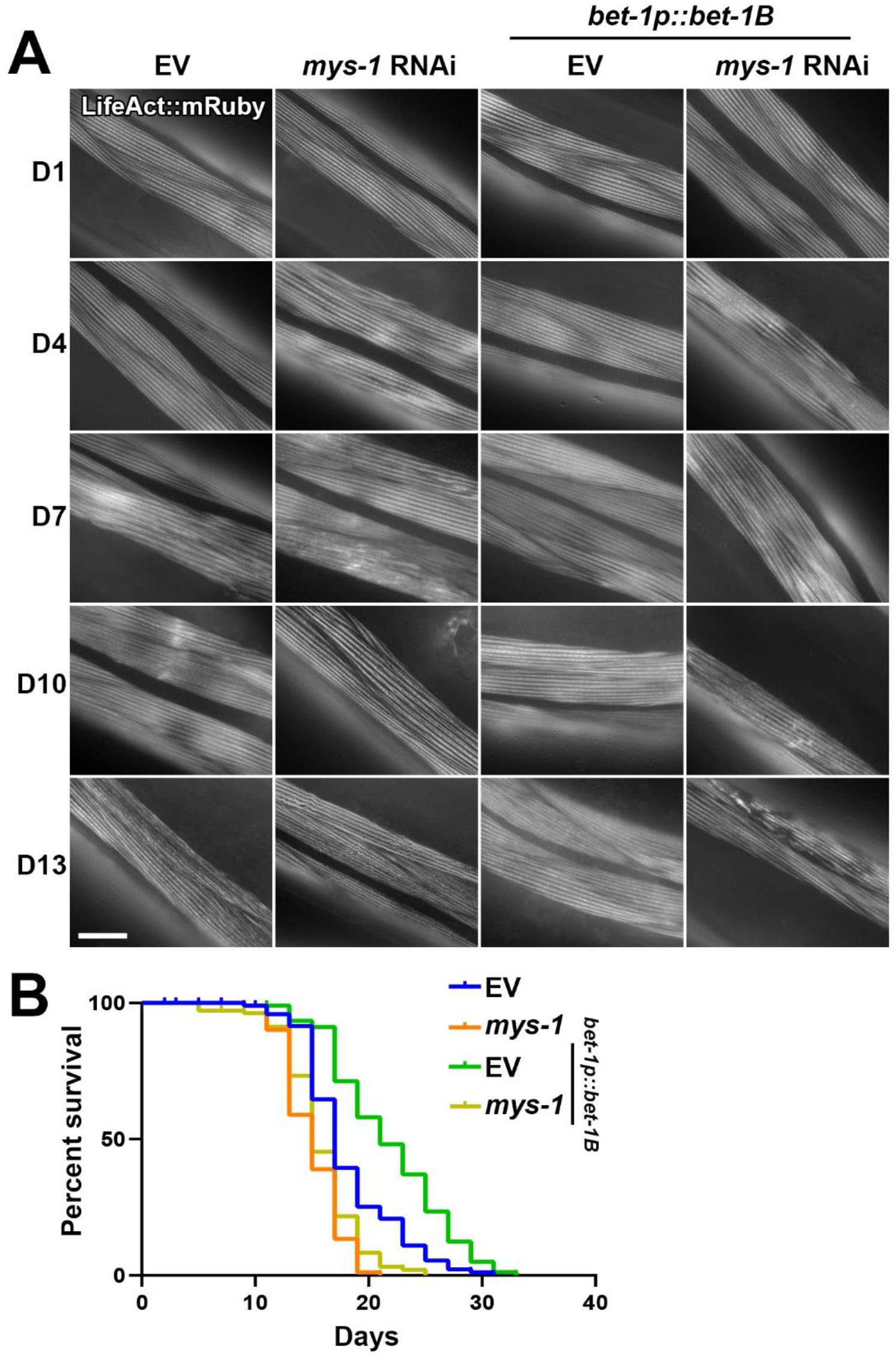
*mys-1* is required for BET-1 mediated effects on actin and lifespan. **(A)** Representative fluorescent images of adult animals expressing Lifeact::mRuby from a muscle-specific promoter *myo-3p*. Wild-type, N2 and *bet-1B* overexpression (*bet-1p::bet-1B*) animals were grown on empty vector (EV) or *mys-1* RNAi from hatch and imaged at day 1, 4, 7, 10, and 13 of adulthood. Images were captured on a Zeiss AxioObserver.Z1. Scale bar is 10 µm. **(B)** Lifespans of N2 (EV, blue) and *bet-1B* overexpression (*bet-1p::bet-1B*, green) animals grown on EV or *mys-1* RNAi (N2, orange; *bet-1B* overexpression, yellow) from hatch. See **Table S3-4** for lifespan statistics.

**Fig. S5.**
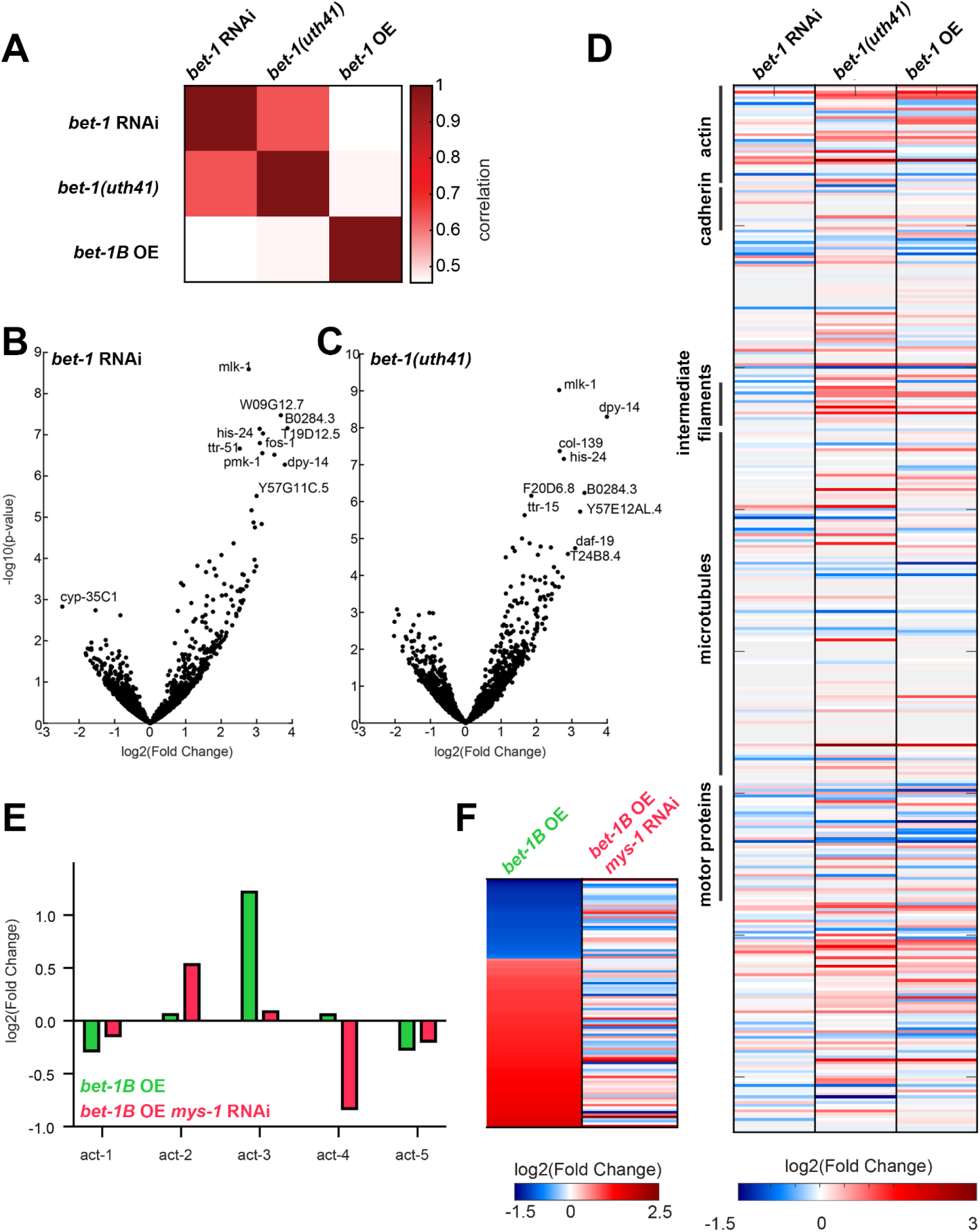
Gene expression analysis of bet-1 knockdown, knockout, and over-expression. **(A)** Pearson’s correlation of log2(fold changes) in gene expression between the different datasets, normalized by N2 wild-type control. Gene expression changes, as in Figure 3A, for *bet-1* RNAi **(B)** or *bet-1* CRISPR mutant **(C)**. **(D)** Gene expression changes for all cytoskeleton genes as annotated in WormCat (Holdorf et al., 2020): actin function; cadherin; centrosome; claudin; innexin; integrin; intermediate filament protein; microtubule; motor protein; other. **(E-F)** Changes in differentially expressed genes is dependent on *mys-1*: Gene expression of *bet-1B* over-expressing worms subjected to *mys-1* RNAi was compared to a *mys-1* RNAi only baseline control. The five actin genes are plotted in **(E)**, and all differentially expressed genes in the *bet-1B* over-expressing worms are plotted in **(F)**. See **Table S5** for all differentially expressed genes.

**Table S1. Enriched genes from cytochalasin screen.**

**Table S2. Depleted genes from cytochalasin screen.**

**Table S3. Statistics for lifespans.**

**Table S4. Replicates for lifespans.**

**Table S5. Differentially expressed genes from RNA-seq analysis.**

**Table S6. Raw sgRNA counts from cytochalasin screen.**

